# Chaperone structure is not a sufficient determinant for the hierarchy of substrate secretion in bacterial type III secretion systems

**DOI:** 10.64898/2026.08.13.744596

**Authors:** Sara V. Pais, Pauline Fauser, Sarah Schroth, Joe Joiner, Elsa Poncet, Sophie Schminke, Marcus Hartmann, Samuel Wagner

**Affiliations:** Interfaculty Institute of Microbiology and Infection Medicine (IMIT), Section of Cellular and Molecular Microbiology, University of Tübingen, Tübingen, Germany; Max Planck Institute for Biology Tübingen, Department of Protein Evolution, Tübingen, Germany; Interfaculty Institute of Biochemistry, University of Tübingen, Tübingen, Germany; Excellence Cluster “Controlling Microbes to Fight Infections” (CMFI), Tübingen, Germany; German Center for Infection Research (DZIF), partner-site Tübingen, Tübingen, Germany

## Abstract

Functional type III secretion in Gram negative bacteria relies on precise substrate targeting and a strict order of secretion with early, intermediate, and late substrates. Type III secretion chaperones facilitate these processes by maintaining substrates in a partially unfolded, secretion-competent state and serving as order-specific targeting factors. Early needle filament assembling substrates are chaperoned by none or class III chaperones, intermediate translocator-type substrates by class II and late effector-type substrates by class I chaperones. In case of hydrophobic transmembrane effectors, chaperones may also serve to prevent erroneous mistargeting of these substrates to the bacterial inner membrane. Here, we characterized the *Salmonella* transmembrane effectors SseF and SseG and their chaperone SscB, encoded in the operon *sscB*-*sseF*-*sseG*, in order to gain a deeper understanding of the underlying molecular requirements of targeting of this special class of substrates. We show that the gene linkage of SscB and SseF is critical for these proteins’ stability and SseF secretion. Counterintuitively, SscB revealed to feature a class II chaperone structure with a class I chaperone function. Likewise, SseF and SseG harbour conserved, translocator-like chaperone-binding motifs (PXI/LXXP) but were secreted as late substrates, independent of the gatekeeper protein SsaL. These findings challenge the current chaperone classification and our understanding of the molecular basis of the hierarchy of substrate secretion. They show that chaperone structure is not a sufficient molecular determinant for the correct order of substrate secretion.

## Introduction

Type III secretion systems (T3SS) are highly specialized protein transport machines, widespread among Gram-negative bacteria and central to bacterial pathogenicity. T3SSs consist of a multiprotein needle-like structure, known as the injectisome, which spans bacterial and host cell membranes. The injectisome forms a channel that directly delivers bacterial proteins into the host cell, where they manipulate various cellular functions (Pais et al., 2023). The secreted proteins, also known as type III secretion (T3S) substrates, include extracellular components of the injectisome, i.e., needle subunits and translocators, as well as effectors. Proteins destined for type III secretion must be targeted to the membrane-embedded injectisome and secreted in a strictly hierarchical order. This process relies on non-cleavable secretion signals, typically located within the first 20 to 40 amino acids of the N-terminus of the T3S substrates (Lloyd et al., 2001; Rüssmann et al., 2002; Sory et al., 1995) that can be recognised across T3SSs from different bacteria (Rosqvist et al., 1995). In addition, many substrates contain a chaperone-binding domain (CBD) through which they interact with specific T3S chaperones.

T3S chaperones maintain substrates in a secretion-competent state by stabilizing and keeping them in a partially unfolded state, a prerequisite for secretion (Stebbins and Galán, 2001; Tucker and Galán, 2000). They serve as targeting factors by interacting with various components of the injectisome, such as the cytosolic sorting platform components SctKQL, the ATPase SctN, the export gate protein SctV (Lara-Tejero et al., 2011), and the gatekeeper SctU (Archuleta and Spiller, 2014). Depending on the class of substrate they bind, T3S chaperones are divided into distinct classes that exhibit common structural features. Class I chaperones bind effectors (late substrates), class II chaperones bind translocators (intermediate substrates), and class III chaperones bind needle subunits (early substrates), thereby reflecting the hierarchy of secretion (Parsot et al., 2003)10/08/2026 12:45:00.

An additional layer of complexity arises from T3S substrates that contain highly hydrophobic transmembrane segments, which can interfere with their targeting to the injectisome. These segments are prone to be recognised co-translationally by the signal recognition particle (SRP), potentially leading to their erroneous insertion into the bacterial inner membrane via the Sec pathway. By binding and masking the transmembrane segments, T3SS chaperones can outcompete the SRP and ensure correct targeting to the injectisome and efficient secretion (Krampen et al., 2018). Transmembrane T3SS substrates include both translocators and effectors (Godlee and Holden, 2023). All T3SSs have at least one pair of translocators together with a dedicated class II chaperone. In contrast, effectors containing transmembrane segments (transmembrane effectors) are present in many bacteria, yet few cognate chaperones have been described; the known pairs include *Salmonella* SscB/SseF (Dai and Zhou, 2004) and SrcA/SteD (Godlee et al., 2019), *Edwardsiella* EseB/EseG (Xie et al., 2010), *E. coli* CesT/Tir (Elliott et al., 1999), and *Vibrio* VecA/VopQ (Iimori et al., 2025). Although shown to aid substrate secretion in early studies (Wattiau and Cornells, 1993), the precise role of T3S chaperones remains unclear, mainly due to the transient and dynamic nature of the secretion process. Secretion of transmembrane T3S substrates adds further complexity to the process, as these substrates are more prone to mistargeting and degradation, thus requiring a tighter regulation.

To investigate the events preceding secretion of transmembrane effectors, we focused on SseF and SseG and the T3S chaperone SscB, encoded in the same operon (*sscB-sseF-sseG*) within the *Salmonella* pathogenicity island 2 (SPI-2). SseF and SseG are among the 8 core effectors conserved across *Salmonella enterica* serovars (Jennings et al., 2017), and are delivered into the host cell by T3SS-2, one of two T3SSs encoded by *Salmonella*. Within the host cell, SseF and SseG interact with each other and cooperate to position the *Salmonella*-containing vacuole (SCV) in proximity to the Golgi, enabling nutrient acquisition and successful intracellular replication (Deiwick et al., 2006; Feng et al., 2018; Hensel et al., 1998; Kuhle and Hensel, 2002; Yu et al., 2016). SscB interacts with SseF through its CBD and first transmembrane segment (Dai & Zhou, 2004; Krampen et al., 2018), an interaction essential for SseF stability (Dai & Zhou, 2004) and correct targeting for secretion (Krampen et al., 2018). We hypothesize that SscB must rapidly interact with SseF upon translation, and that gene location may be critical to spatiotemporally connect the translation of SscB and SseF, thereby facilitating the formation of the SscB/SseF complex and ensuring confident secretion of SseF. To test these hypotheses, we examined genomic organisation, protein stability, chaperone/effector interactions and secretion using a combination of biophysical and biochemical approaches, as well as secretion and injection assays based on the split-NanoLuc luciferase reporter. We found that the specific gene location within the operon is required for efficient secretion of SseF, enabling the formation of the SscB/SseF complex and stabilizing both proteins. For SseG, interaction with SscB appears to be important for secretion but not for stability. Characterisation of the chaperone/effector complexes revealed that SscB exhibits structural features and a mode of interaction typical of class II chaperones despite SseF and SseG being bona fide effectors rather than translocators. Overall, these results demonstrate that the events that precede transmembrane substrate secretion are interdependent and that class II chaperones can also bind and aid secretion of transmembrane effectors, challenging the current paradigm of T3S chaperone classification.

## Results

### The genomic location of *sscB* directly upstream *sseF* facilitates secretion of SseF

To study the events leading to the secretion of T3S effectors through the T3SS-2 of *Salmonella*, we adapted the split-NanoLuc secretion assay previously described to assess secretion through T3SS-1 (Westerhausen et al., 2020). *Salmonella* strains expressing the effector with a C-terminal fusion to the HiBiT tag (SseF^HiBiT^) were grown under conditions that induce SPI-2 gene expression and promote the secretion of substrates via T3SS-2 (Löber et al., 2006). Detection of the secreted proteins was achieved by the addition of purified LgBiT and the NanoLuc substrate furimazine followed by luminometry. This assay allowed for the analysis of bacterial cultures rather than supernatant fractions. This is key for the detection of SseF secretion, as this protein is detected not only in the supernatant, but also at the bacterial surface (Hansen-Wester et al., 2002) (Supp. Text 1, Fig. S1 and S2).

Given that binding of the T3S chaperone SscB to the transmembrane effector SseF may need to occur rapidly to prevent mistargeting, we hypothesised that the location of the *sscB* gene upstream of *sseF* is required for efficient type III secretion of SseF. To assess this, secretion of SseF was compared between proteins expressed from the original and reverse gene order. We used *Salmonella* strains that do not express SscB (*sscB*), harbouring plasmids that expressed SscB and SseF (pSseF, pSscB-SseF and pSseF-SscB) (Fig. 1A). For better comparison, an optimised Shine-Dalgarno sequence was added in front of each gene in all plasmids, a strategy used throughout this work. The analysis of SseF secretion showed that changing the gene order (pSseF-SscB) led to a reduction in the secretion of SseF to 62 ± 12 % of the original gene order (pSscB-SseF) (Fig. 1C), despite similar amounts of protein being detected in both *sscB* and *sscB ssaV* strains (Fig. 1B). No secretion was observed in the absence of the chaperone (pSseF) and in T3S-deficient strains (*sscB ssaV*) (Fig. 1C). Further, we analysed the injection of SseF during infection using the same *Salmonella* strains. We used the split-NanoLuc assay to assess the injection via T3SS-2 into host cells (Fig. S1)(Boudrioua et al., 2025). The change in gene order (pSseF-SscB) led to a drastic decrease in translocated SseF, dropping to 5 ± 3 % compared to the original gene order (pSscB-SseF) (Fig. 1D). Similar to the secretion results, injection of SseF was neither detected in the strain expressing only SseF (*sscB*; pSseF), nor in any of the T3S-deficient strains (*sscB ssaV*) (Fig. 1D). *Salmonella sscB* and *sseF* mutants show a reduced intracellular replication compared to the wild-type strain (Kuhle and Hensel, 2002). Therefore, we reasoned that the different effects of the reversed gene order on secretion and injection may be the consequence of a cumulative effect of the decreased secretion/injection of SseF and of the inability of *Salmonella* to sustain infection in the host cells in the absence of SseF. To confirm this, HeLa-LgBiT cells were infected with *Salmonella* and after 16 h, colony-forming units (CFU) were enumerated. Indeed, the change in gene order (pSseF-pSscB) led to a more than four-fold reduction in CFU count when compared with the original gene order (pSscB-SseF) (Fig. 1E).

**Figure 1.**
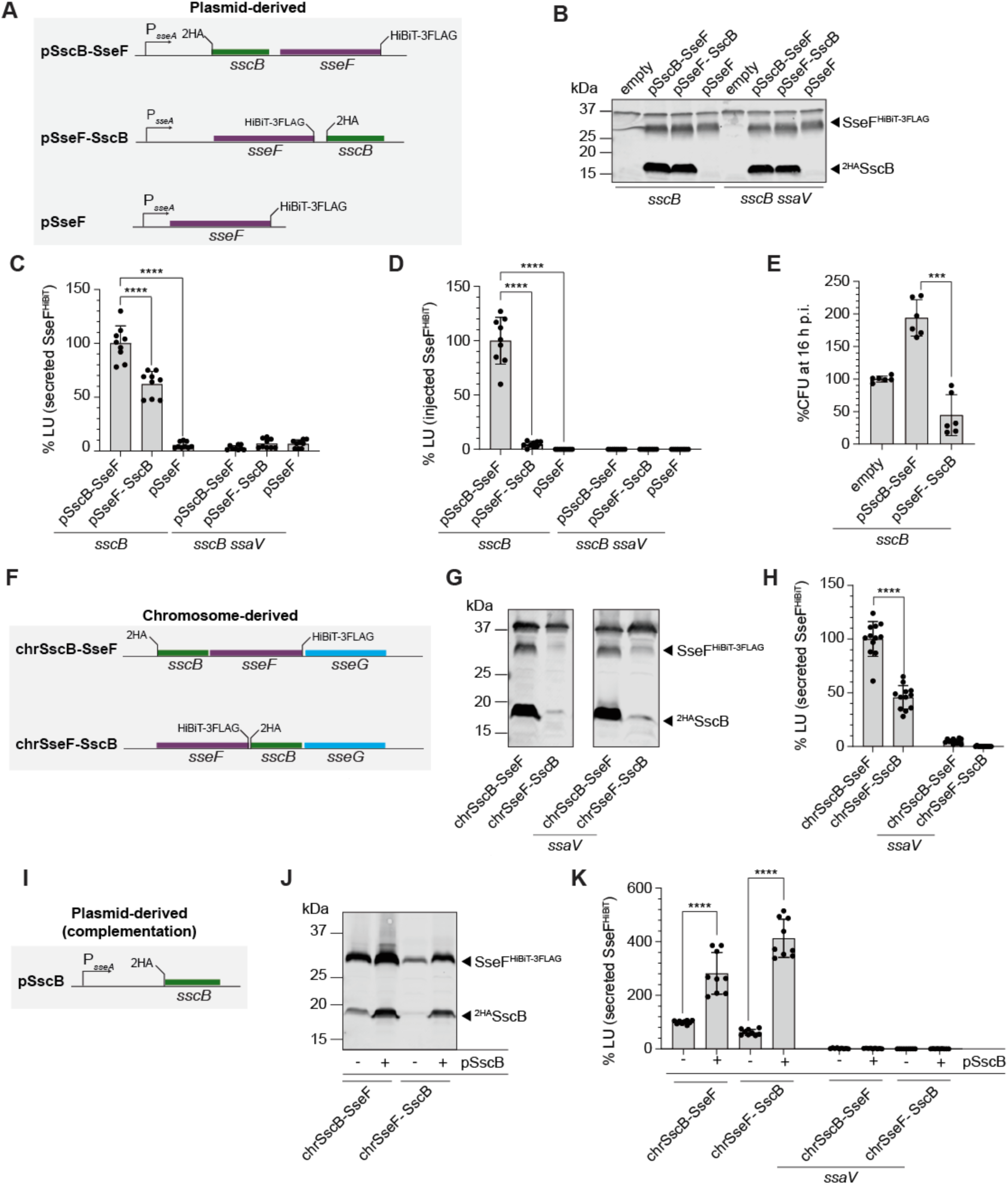
The genomic location of *sseF* downstream of *sscB* in the genome is essential for SseF secretion and injection. **(A)** The role of the genomic organisation was analysed using plasmids encoding SscB and SseF, with the depicted gene location, under the control of the endogenous promoter (P_sseA_). **(B)** and **(C)** T3S-proficient (*sscB*) and T3S-deficient (*sscB ssaV*) *Salmonella* strains were grown under SPI-2-inducing and secreting conditions **(B)** Whole-cell extracts were analysed by SDS-PAGE and immunoblotted using anti-HA and anti-FLAG antibodies. **(C)** Bacterial cultures were collected, and secretion was analysed using the split-NanoLuc secretion assay. The secretion of plasmid-derived SseF^HiBiT^ was analysed by luminometry. The luminescence signal was compared to the T3S-proficient strain (*sscB*) expressing SscB and SseF (pSscB-SseF). **(D)** and **(E)** HeLa cells constitutively expressing LgBiT were infected with the indicated *Salmonella* strains for 16 h. **(D)** SseF^HiBiT^ injected into host cells was analysed by measuring the luminescence signal (LU) at 16 h post-infection (p.i.). The luminescence signal was compared to the T3S-proficient strain (*sscB*) expressing SscB and SseF from the original gene order (pSscB-SseF). **(E)** At 16 h p.i., infected cells were lysed and plated on agar media to enumerate the colony-forming units (CFU). The CFU were compared to the *sscB* mutant strain without a plasmid. **(F)** The role of the genomic organisation was analysed using *Salmonella* strains encoding tagged SscB and SseF within the chromosome in the depicted gene location (chrSscB-SseF, chrSseF-SscB). **(G)** and **(H)** T3S-proficient and T3S-deficient (*ssaV*) *Salmonella* strains were grown under SPI-2-inducing and secreting conditions. **(G)** Whole-cell extracts were analysed by SDS-PAGE and immunoblot using anti-HA and anti-FLAG antibodies. **(H)** Bacterial cultures were collected, and secretion was analysed using the split-NanoLuc secretion assay. The secretion of chromosome-derived SseF^HiBiT^ was analysed by luminometry (LU). The luminescence signal was compared to the T3S-proficient strain expressing SscB and SseF in the original gene order (chrSscB-SseF). **(I)** The *Salmonella* strains encoding tagged SscB and SseF within the chromosome, as depicted in (F), were complemented with a plasmid encoding SscB under the control of its endogenous promoter (P_sseA_). **(J)** T3S-proficient *Salmonella* strains encoding tagged SscB and SseF within the chromosome (chrSscB-SseF, chrSseF-SscB) were complemented (+) or not (-) with plasmid-derived SscB (pSscB). Bacteria were grown under SPI-2-inducing and secreting conditions for 6 h, then harvested and analysed by SDS-PAGE and immunoblot using anti-HA and anti-FLAG antibodies. **(K)** Bacterial cultures were collected, and secretion was analysed using the split-NanoLuc secretion assay. The secretion of chromosome-derived SseF^HiBiT^ was analysed by luminometry (LU). The luminescence signal was compared to the T3S-proficient strain expressing SscB and SseF in the original gene order (chrSscB-SseF). Statistical significance is indicated as follows: **** P<0.0001; *** P<0.001, further description of the statistical analysis can be found in Supp. Table 3.

Next, a similar analysis was performed using *Salmonella* mutants with a reversed gene order on the chromosome (chrSscB-SseF and chrSseF-SscB) (Fig. 1F). SDS-PAGE and immunoblot analysis revealed that changing the gene order (chrSseF-SscB) decreased the accumulation levels of both SscB and SseF inside bacteria as compared to the original gene order (chrSscB-SseF) (Fig. 1G), an effect that was not observed upon plasmid-based complementation (Fig. 1B), possibly due to overexpression of the respective proteins. Secretion of SseF decreased to 46 ± 11% upon change of gene order (chrSscF-SscB) (Fig. 1H), but complementation of the chromosomal mutants (chrSscB-SseF and chrSseF-SscB, respectively) with plasmid-derived SscB (pSscB) (Fig. 1I) resulted in an overall increase in SseF’s protein levels (Fig. 1J). The complemented strains displayed a strong increase in the secretion of SseF irrespective of the gene order (Fig. 1K). To rule out a possible regulatory function of SscB in overall secretion, we evaluated the effect of overexpressing SscB in *trans* on the secretion of PipB2, a T3S effector secreted through T3SS-2. Chromosome-derived PipB2 was expressed (Fig. S3A) and secreted (Fig. S3B) in similar levels both with and without plasmid-derived SscB (pSscB).

Altogether, our results showed that changing the order of *sscB* and *sseF* reduced the secretion/injection of SseF and decreased *Salmonella*’s fitness in the host cell. This demonstrates that the gene order is an important requirement for the efficient secretion of SseF and *Salmonella* pathogenicity. However, overexpression of SscB in *trans* was able to override the detrimental effects of a change in gene order. Together, this indicates that translation of *sscB* upstream *sseF* and the amount of available SscB are the limiting factors for stability and secretion of SseF.

### SscB is a type III secretion chaperone of SseG by aiding its secretion

The *sscB-sseF-sseG* operon encodes a second transmembrane effector, SseG, which is hypothesized to also be chaperoned by SscB. Previous attempts to assess whether SscB aided secretion of SseG were unsuccessful (Dai and Zhou, 2004), possibly due to low sensitivity of the methodologies employed. Using the split-NanoLuc secretion assay, we analysed *Salmonella* strains expressing the plasmid-derived effectors alone (SseF and SseG) or in combination with SscB (SscB + SseF and SscB + SseG). Secretion of both SseF and SseG was detected only in the presence of SscB (Fig. 2A). SseG was less secreted (11 ± 3 %) than SseF (98 ± 5 %), despite similar amounts of the two effectors detected in the whole-cell extracts (Fig. 2B). SseG was expressed even in the absence of SscB, although no secretion could be detected under these conditions, while SscB was more readily detected when co-expressed with SseF rather than with SseG (Fig. 2B). Together, these results confirm that SscB is not only a type III secretion chaperone of SseF but also of SseG, although it supports secretion of SseF more effectively than that of SseG.

**Figure 2.**
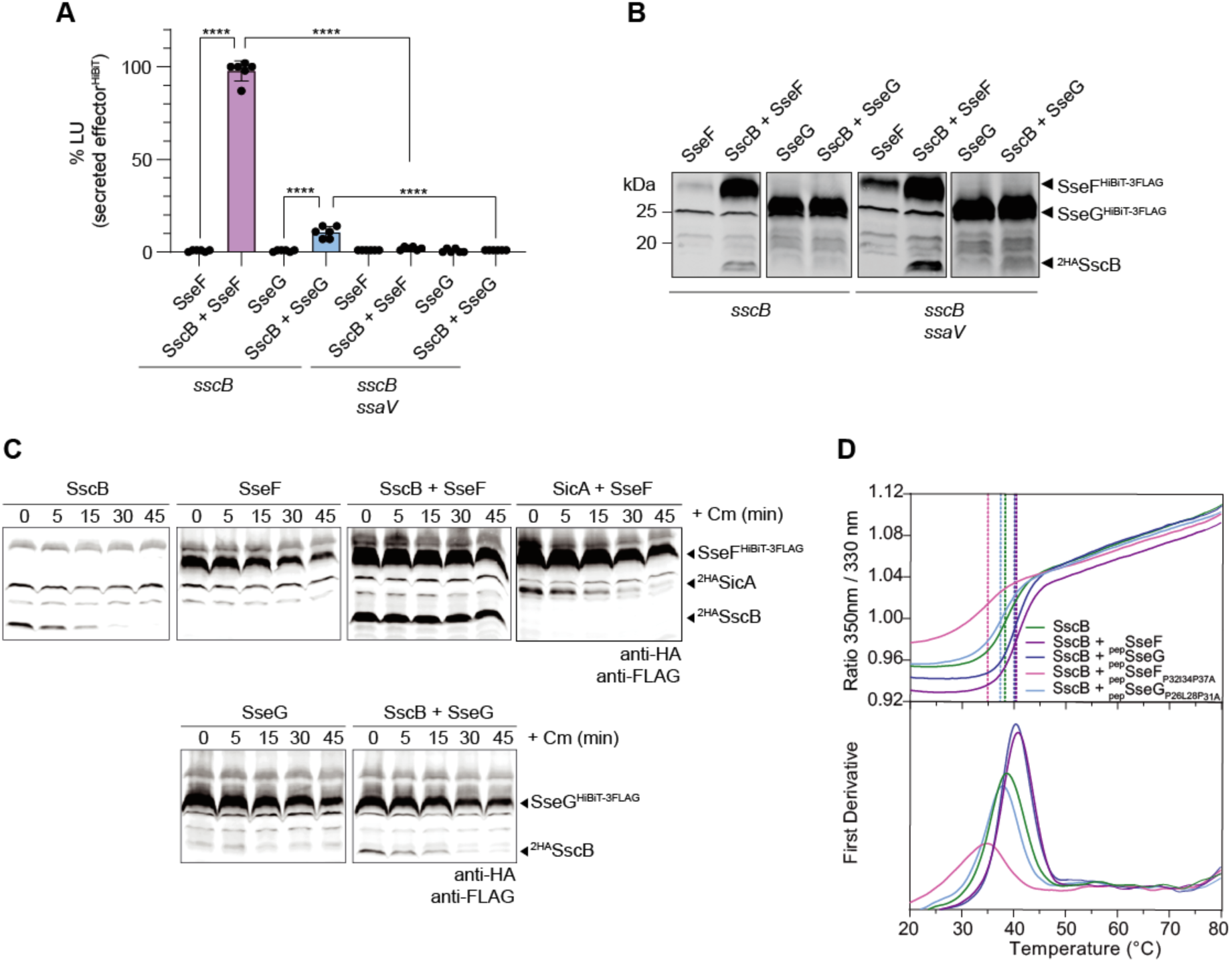
SscB aids the secretion of both SseF and SseG, and is itself stabilized by SseF. *Salmonella* T3S-proficient (*sscB*) or T3S-deficient (*sscB ssaV*) strains were grown under SPI-2-inducing and secreting conditions. These strains harboured plasmids expressing SseF or SseG alone or with SscB (SscB + SseF and SscB + SseG), under the control of a rhamnose-inducible promoter. **(A)** Bacterial cultures were collected, and secretion was analysed using the split-NanoLuc T3S assay. The secretion of SseF^HiBiT^ or SseG^HiBiT^ was analysed by luminometry. The luminescence signal was compared to the T3S-proficient strain (*sscB*) expressing both SscB and SseF (SscB + SseF). **(B)** Whole-cell extracts were analysed by SDS-PAGE and immunoblot using anti-HA and anti-FLAG antibodies. **(C)** Protein stability was analysed in T3S-deficient (*sscB ssaV*) *Salmonella* strains expressing ^2HA^chaperone/effector^HiBiT-3FLAG^ pairs from a rhamnose-inducible (P_rha_) promoter. Bacteria were grown under SPI-2-inducing conditions, and protein expression was induced when OD_600_ reached 0.5. After 3 h of induction, chloramphenicol (Cm) was added, and samples were harvested every 15 min. Whole-cell extracts were analysed by SDS-PAGE immunoblot using antibodies against HA and FLAG. Time zero (0 min) corresponds to time before the addition of chloramphenicol. **(D)** NanoDSF unfolding profiles of purified SscB when incubated with SseF and SseG peptides containing the wild-type and triple alanine mutations in the chaperone-binding domain (CBD) motif. The SscB melting temperature calculated for all the samples is indicated by the dashed vertical lines. Statistical significance is indicated as follows: **** P<0.0001, further description of the statistical analysis can be found in Supp. Table 3.

### SscB is stabilised by SseF but not by SseG

The differences in protein levels of SscB, SseF and SseG (Fig. 1G and 2B) led us to investigate the stability of these proteins. T3S-deficient *Salmonella* strains (*sscB ssaV)* expressing the proteins individually (SscB, SseF, SseG) or as a chaperone/effector pair (SscB + SseF, or SscB + SseG) were grown under SPI-2 inducing conditions. After induction of protein expression for 3 h, chloramphenicol was added to halt protein synthesis, and protein stability was analysed over time (Fig. 2C and S4). In agreement with previous reports, SseF was stabilized when co-expressed with SscB (Dai and Zhou, 2004). Reciprocally, SscB was quickly degraded when expressed alone but stabilized upon co-expression of SseF (Fig. 2C and S4), demonstrating that SscB and SseF stabilize each other. SseG remained more stable than SseF in the absence of SscB, while SscB did not maintain the same level of stability in the presence of SseG as it did with SseF (Fig. 2C and S4). Note that although the degradation profile shows similar half-lives (Fig. S4), the amount of protein at time zero should also be considered, as it reflects different turnover rates during the induction of protein expression. Next, the melting temperature (T_m_) of purified SscB was measured using nano differential scanning fluorimetry (NanoDSF) (Fig. 2D). SscB alone had a T_m_ of 38.62 °C, consistent with the observations that SscB alone was unstable when expressed in *Salmonella* at 37 °C (Fig. 2C). The addition of peptides containing only the chaperone-binding domains (CBD) of either SseF or SseG (further characterised in the following sections) led to an increase in the melting temperature of SscB by approximately 2 °C (_pep_SseF, T_m_ = 40.79 °C and _pep_SseG, T_m_ = 40.38 °C) (Fig. 2D), whereas the differences between the melting temperatures of SseF and SseG peptides were only 0.41 °C. Taken all together, these results showed that SscB and SseF require each other for stability, in contrast to SseG, which is more stable by itself and unable to stabilise SscB. Moreover, the chaperone-binding domains of SseF and SseG alone were sufficient to stabilize SscB *in vitro*.

### Mapping the interactions of SscB/SseF and SscB/SseG complexes using in vivo photocrosslinking

While SscB was shown to interact with the chaperone-binding domain and the first transmembrane segment of SseF (Dai and Zhou, 2004; Krampen et al., 2018), the interaction between SscB and SseG has not been reported previously. To map interaction sites within the chaperone/effector complexes, we used site-specific *in vivo* photocrosslinking. This method relies on the incorporation of the UV-reactive unnatural amino acid *para*-benzoyl-phenylalanine (*p*Bpa) at selected positions, introduced by mutagenesis to an amber stop codon (denoted by X). Upon UV irradiation, *p*Bpa forms covalent crosslinks with neighbouring proteins. Briefly, *Salmonella* strains encoding chaperone or effector amber mutants (SscB_x_, SseF_x_ and SseG_x_) were grown under SPI-2-inducing conditions, irradiated with UV to induce *p*Bpa-derived crosslinks, and further analysed by SDS-PAGE and immunoblot.

For the SscB/SseF complex (Table S1), faint but crosslinking-specific bands were observed for SscB variants carrying *p*Bpa at residues M3 (SscB_M3X_) and W56 (SscB_W56X_) (Fig. 3A). The crosslink band of SscB_M3X_ was detected by anti-FLAG (SseF^HiBiT-3FLAG^) but not anti-HA antibodies (^2HA^SscB). In contrast, the crosslink band of SscB_W56X_ was only detected with anti-HA antibody. For SseF-*p*Bpa mutant variants (Fig. 3A), stronger crosslinking-specific bands were detected for residues A24X, L69X and V73X with both antibodies, as observed previously (Krampen et al., 2018). A weaker crosslinking-specific band was observed for residue L169X, with both antibodies. The higher molecular mass band at approximately 50 kDa corresponds to the predicted molecular mass of the complex. The lower molecular mass bands, with a difference of less than 10 kDa, may correspond to crosslinks with a N-terminal cleaved form of SscB, which is untagged and therefore not detected by immunoblot.

**Figure 3.**
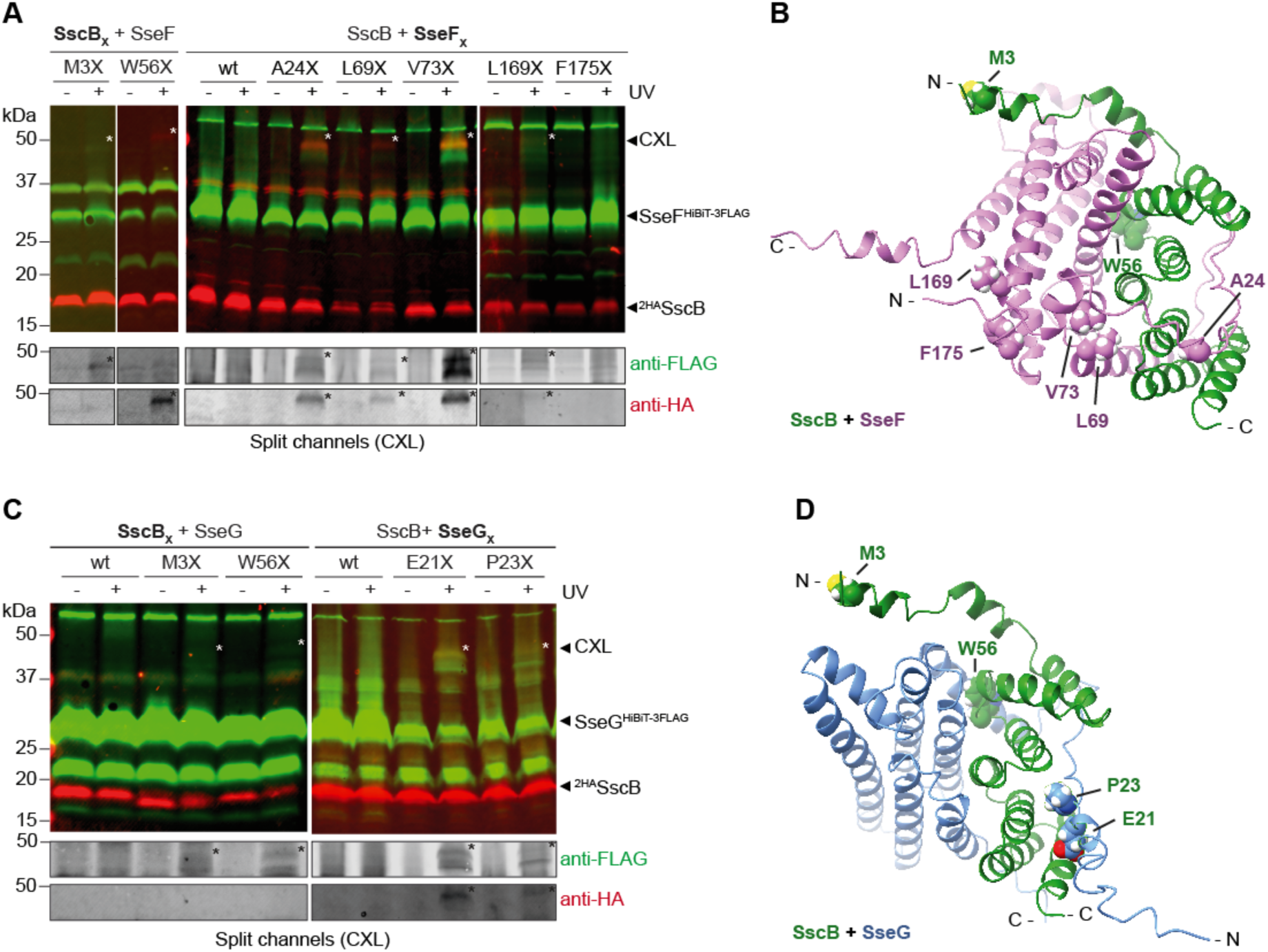
Mapping of interactions in SscB/SseF in SscB/SseG complexes by *in vivo* photocrosslinking. *Salmonella sscB* mutant strains harbouring plasmids co-expressing mutant variants of **(A)** SscB and SseF, or **(C)** SscB and SseG (pBpa incorporation denoted as X), under the control of a rhamnose-inducible promoter, were grown under SPI-2 inducing and secreting conditions. Bacteria were harvested and exposed to UV for 30 min (+UV) or not (-UV), followed by analysis by SDS-PAGE and immunoblot using antibodies against HA (red) and FLAG (green). Crosslinks are indicated as CXL and *. Structures of **(B)** SscB and SseF (green/pink) and of **(D)** SscB and SseG (green/blue) were predicted using Alphafold2 Multimer. The residues mutated for pBpa incorporation are depicted in the structures.

Next, we analyzed whether SscB and SseG form a complex using *in vivo* photocrosslinking. The SscB mutant variants SscB_M3X_ and SscB_W56X_ (Fig. 3C), showed weak but crosslinking-specific bands that were both detected only with anti-FLAG antibodies (SseG^HiBiT-3FLAG^). When we tested several SseG-*p*Bpa mutant variants (Table S2), crosslinking-specific bands at the predicted molecular mass of the complex, 48 kDa, were only detected for residues E21X and P23X, alongside lower molecular mass bands, as observed for SscB/SseF (Fig. 3C).

To further characterize the chaperone-effector interactions, we purified the SscB/SseF complex and performed size-exclusion chromatography coupled with multi-angle light scattering (SEC-MALS), which showed a complex with a molecular mass of 51 ± 2 kDa, indicating a 1:1 stoichiometry (Fig. S5). For SscB/SseG, not enough protein could be purified to perform SEC-MALS. Furthermore, structures of SscB/SseF and SscB/SseG were predicted using Alphafold2 Multimer. SscB in both SscB/SseF and SscB/SseG complexes had a high confidence score (pLDDT > 95), except for the N-terminal region (residues 1 to 11; pLDDT < 70) (Fig. S6). In the prediction of the SscB/SseF complex, SseF had overall low confidence scores, except for the regions corresponding to the chaperone-binding domain (residue 29 to 42) and the first transmembrane segment (residue 45 to 85), which exhibited the highest confidence scores (pLDDT > 70) (Fig. S6A). In the SscB/SseG complex prediction, SseG only had a high confidence score (pLDDT > 70) between residues 22 and 32 (Fig. S6B).

Overall, our data showed that the N-terminal region of SscB crosslinks with both SseF and SseG, suggesting a conserved mode of interaction. Furthermore, SscB interacts not only with SseF’s chaperone-binding domain and the first transmembrane segment, but also with the second transmembrane segment (L169X). Regarding SseG, interactions were mapped through residues E21 and P23, which are predicted to comprise the chaperone-binding domain, based on comparison of the predicted structures of SscB/SseF and SscB/SseG complexes (Fig. 3B and 3D).

### SscB is structurally similar to class II chaperones of translocators

Based on sequence similarity, SscB is annotated as a “CesD/SycD/LcrH family type III secretion system chaperone” (Cirillo et al., 1998), classifying it as a class II, intermediate substrate-specific chaperone. However, SscB binds and aids secretion of SseF and SseG, bona fide T3S effectors, and should be functionally classified as a class I T3S chaperone (Parsot et al., 2003). To clarify this paradox, we first examined the structural characteristics of SscB. The monomeric structure of SscB predicted by Alphafold2 had a high confidence score (pLDDT > 95), with exception of the N-terminal region (residues 1 - 11; pLDDT < 70) and revealed an α-helical fold (Fig. 4A). This was confirmed by circular dichroism spectroscopy (CD) of purified SscB, as the CD spectrum of SscB, with minima around 208 nm and 222 nm, was compatible with the CD spectrum of an α-helical protein (Fig. 4B). Thus, SscB does not adopt the typical ɑ/β sandwich fold of class I chaperones, but instead an ɑ-helical fold, characteristic of class II chaperones. In solution, purified SscB forms dimers, with SEC-MALS analysis revealing a molecular mass of 31.1 ± 1.2 kDa, approximately twice the predicted molecular mass of a SscB monomer (16.7 kDa). Further supporting its uniqueness among chaperones of effectors, the predicted structure of SscB is structurally homologous to the solved crystal structures of SycD from *Yersinia*, PcrH from *Pseudomonas*, IpgG from *Shigella*, and AcrH from *Aeromonas*, all class II T3S chaperones binding translocators (Fig. 4D). Within the SscB/SseF complex, SscB is predicted to have a curved structure in which the concave inner surface forms a binding pocket that accommodates the chaperone-binding domain of SseF (Fig 3B, 4D and 4E). The convex outer surface forms an interface with the first transmembrane segment of SseF (Fig. 3B), reminiscent of the mode of interaction previously described for class II chaperone/translocator (Nguyen et al., 2015).

**Figure 4.**
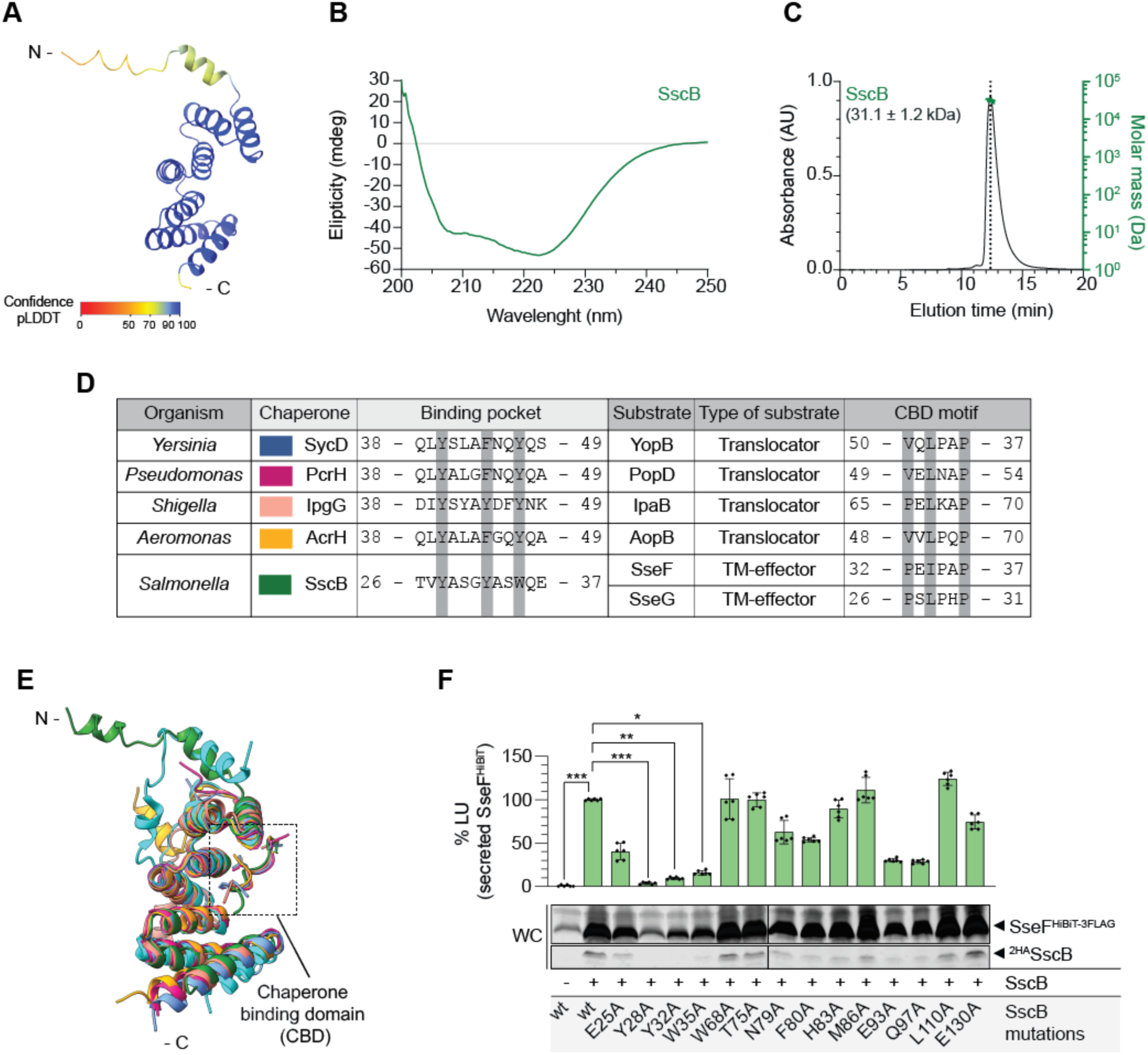
Characterisation of the T3S chaperone SscB. **(A)** Predicted structure of SscB using Alphafold2. The colour gradient indicates the confidence score of the prediction (pLDDT). **(B)** Circular dichroism (CD) spectrum of purified, untagged SscB. **(C)** SEC-MALS (size-exclusion chromatography coupled with multi-angle light scattering) chromatogram of purified, untagged SscB. **(D)** Comparison of class II secretion chaperones and their respective substrates. Highlighted in gray are the conserved sequences of the chaperone-binding pocket (in the chaperone) and the conserved CBD motif (in the substrate). **(E)** Overlap of the structures of class II chaperones in complex with a peptide (inset) containing the chaperone-binding domain (CBD) motif of the respective interacting partner (T3S translocator) from different organisms. SycD from *Yersinia* (blue; PDB 4AM9), PcrH from *Pseudomonas* (pink; PDB 2XCB), IpgG from *Shigella* (salmon; PDB 3GZ2), AcrH from *Aeromonas* (yellow; PDB 3WXX), and SscB from *Salmonella* (green; Alphafold2). **(F)** T3S-proficient *Salmonella* strains (*sscB*) co-expressing SscB and SseF under a rhamnose-inducible promoter with the depicted SscB alanine mutations were grown under SPI-2-inducing and secreting conditions. Bacterial cultures were collected, and secretion of SseF^HiBiT^ was analysed by luminometry using the split-NanoLuc T3S assay. The luminescence signal was compared to the T3S-proficient strain (*sscB*) expressing both SscB and SseF wild-type. Whole-cell extracts (WC) were collected and analysed by SDS-PAGE and immunoblot using anti-HA and anti-FLAG antibodies. Statistical significance is indicated as follows: * P<0.05, ** P<0.01, *** P<0.001, further description of the statistical analysis can be found in Supp. Table 3.

We then sought to identify the residues of SscB critical for binding and type III secretion of SseF. We performed alanine scanning mutagenesis of SscB in residues predicted to locate at the interface of the chaperone-binding domain and of the transmembrane segments of SseF (Fig. 4F and 5C). *Salmonella* strains co-expressing, from a plasmid, SscB alanine mutants together with SseF were grown in SPI-2 inducing and secreting conditions, followed by assessment of secretion. Expression of the alanine mutants SscB_Y28A_, SscB_Y32A_ and SscB_W35A_ drastically reduced the secretion of SseF to 4 ± 1 %, 8 ± 1 % and 16 ± 3 %, respectively, compared to wild-type SscB (Fig. 4F). Residues Y28, Y32, and W35 of SscB are predicted to form the interface with the CBD of SseF (Fig. 5C, 5D and 5E) and SseG (Fig. 5G and 5H).

**Figure 5:**
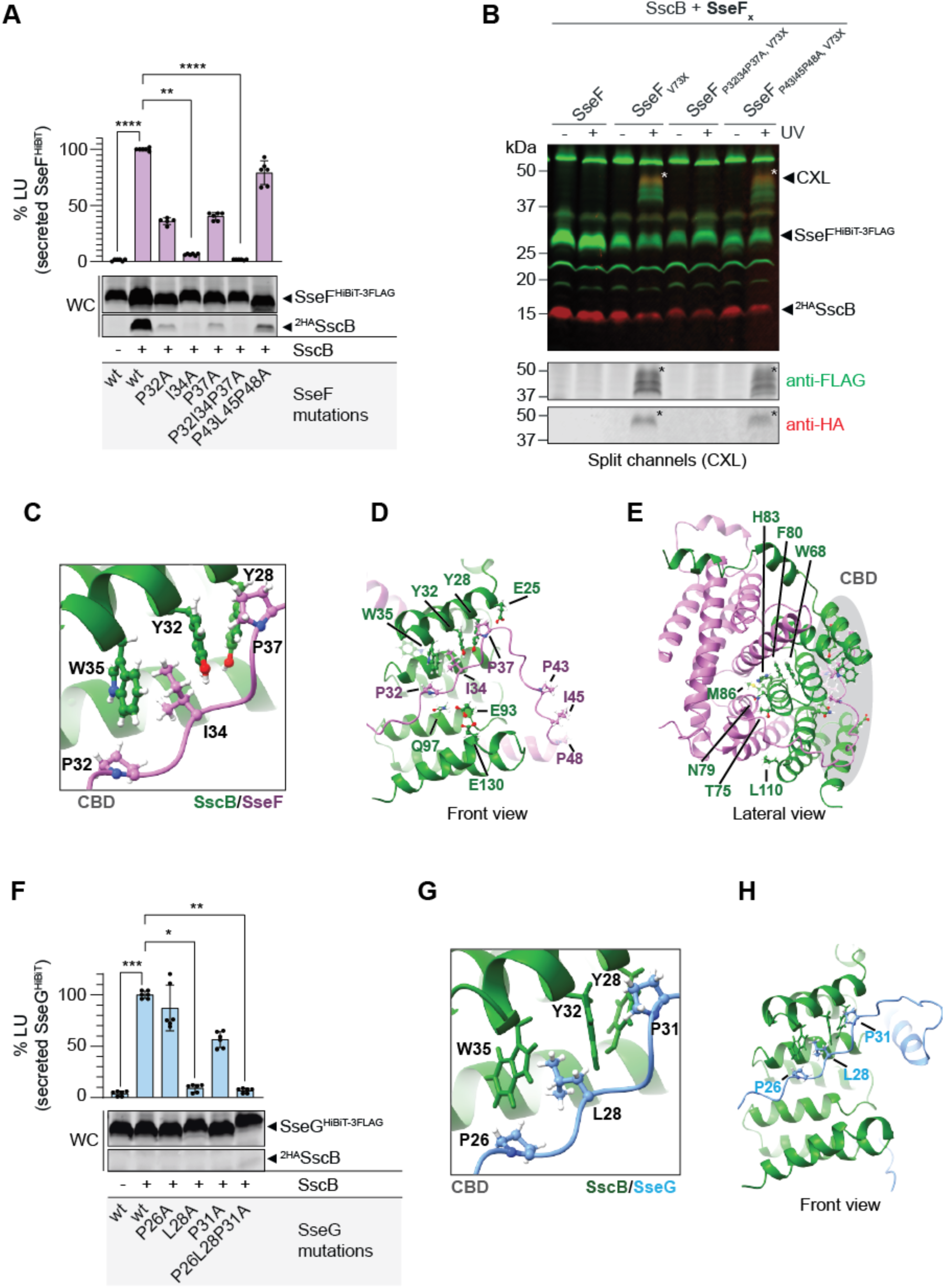
The effectors SseF and SseG harbour a conserved N-terminal chaperone-binding motif. **(A)** T3S-proficient *Salmonella* strains (*sscB*) harbouring plasmids expressing SseF wild-type or with alanine mutation in the chaperone-binding domain were grown under SPI-2-inducing and secreting conditions. Whole-cell extracts were collected and analysed by SDS-PAGE and immunoblot using anti-HA and anti-FLAG antibodies. Bacterial cultures were collected and secretion was analysed using the split-NanoLuc T3S assay. The secretion of plasmid-derived SseF^HiBiT^ was analysed by luminometry. The luminescence signal was compared to the T3S-proficient strain (*sscB*) expressing both wild-type SscB and SseF. **(B)** *In vivo* photocrosslinking was used to analyse the interaction between SscB and SseF. *Salmonella* strains expressed plasmid-derived SscB and SseF (SscB + SseF), where the residue V73 of SseF was mutated for pBpa incorporation (V73X). SseF also harboured triple alanine mutations in the putative CBD motifs. Bacteria were grown under SPI-2 inducing and secreting conditions. Bacteria were harvested and exposed to UV for 30 min (+UV) or not (-UV) and then analysed by SDS-PAGE and immunoblot using antibodies against HA (red) and FLAG (green). Crosslinked SseF with SscB is indicated as CXL and *. **(C), (D)** and **(E)** Predicted structure of the SscB/SseF complex. **(C)** Detailed view of alanine-mutated residues in SscB and SseF. **(D)** Overall structure depicting the chaperone**-**binding domain pocket interface, and **(E)** the transmembrane segment binding interface. **(F)** T3S-proficient *Salmonella* strains (*sscB*) expressing SseG, wild-type or with alanine mutation in the chaperone-binding domain, were grown under SPI-2-inducing and secreting conditions. Whole-cell extracts were collected and analysed by SDS-PAGE and immunoblot using anti-HA and anti-FLAG antibodies. Bacterial cultures were collected, and secretion was analysed using the split-NanoLuc T3S assay. The secretion of plasmid-derived SseG^HiBiT^ was analysed by luminometry. The luminescence signal was compared to the T3S-proficient strain (*sscB*) expressing both wild-type SscB and SseG. **(G)** and **(H)** Predicted structure of the SscB/SseG complex. **(G)** Detailed view of alanine-mutated residues in SscB and SseG. **(H)** Overall structure depicting the chaperone**-**binding domain pocket interface, Statistical significance is indicated as follows: * P<0.05, ** P<0.01, *** P<0.001, ****P<0.0001, further description of the statistical analysis can be found in Supp. Table 3.

Their substitution to alanine may decrease effector binding and result in decreased SseF secretion. Reduced accumulation levels of SscB and SseF in these SscB mutants also supports the notion that the stability of SscB and SseF is mutually interdependent (Fig. 4F), further contributing to a reduced secretion of SseF. Together, our results showed that SscB, despite binding bona fide effectors, structurally resembles a class II chaperone of translocators.

### The effectors SseF and SseG contain a P/VXLXXP motif conserved in translocators

The chaperone-binding domain of translocators harbours a common motif (P/VXLXXP) with three highly conserved residues (Fig. 4D). Based on the similarity of SscB to class II T3SS chaperones, we expected SseF and SseG to harbour a similar motif. In SseF, we identified two putative motifs: <u>P</u>EIPA<u>P</u> (residues 32 - 37) and <u>P</u>V<u>L</u>LT<u>P</u> (residues 43 - 48). Analysis of *Salmonella* strains expressing SscB together with SseF mutant variants in the conserved residues of the putative motifs (Fig. 5A) revealed a reduction in secretion to 36 ± 3 % and 40 ± 3 % in strains expressing SseF_P32A_ and SseF_P37A_, respectively, compared to wild-type SseF (Fig. 5A). Most remarkable was the reduction of secretion to 6 ± 1 % of strains expressing SseF_I34A_. In addition, the triple mutant SseF_P32I34P37A_ had a reduction in secretion down to 2 ± 0 %, while in the SseF_P43L45P48A_ mutant strain, secretion was 79 ± 11 % of that of wild-type SseF strains. Furthermore, the protein levels of the SseF mutants were similar to those of wild-type SseF (Fig. 5A), in contrast to what was previously observed for the SscB alanine mutants (Fig. 4F). Nonetheless, in strains expressing SseF alanine mutants with lower levels of secretion, the lower amounts of SscB were detected by immunoblot (Fig. 5A), suggesting a disrupted SscB/SseF interaction that resulted in protein instability. To assess whether alanine mutations in SseF disrupted the interaction between SscB and SseF, we used *in vivo* photocrosslinking. A *Salmonella* strain harbouring SscB and SseF with an amber mutation at valine 73 (SscB + SseF_V73X_) was used as a background strain. Valine 73 is located within the first transmembrane segment of SseF and was shown to crosslink with SscB (Fig. 2A) (Krampen et al., 2018). Analysis of the alanine triple mutants of SseF (SseF_P32I34P37A,V73X_ and SseF_P43L4548A,V73X_) showed no crosslinking-specific band for SseF_P32I34P37A,V73X_, while crosslinking-specific bands were detected in both wild-type SseF and SseF_P43L4548A,V73X_ (Fig. 5B). As previously discussed, more than one crosslinked band was observed.

Overlap of the predicted structure of SscB/SseF and SscB/SseG showed that the chaperone-binding pocket of SscB also formed an interface with a putative CBD motif of SseG (<u>P</u>S<u>L</u>PH<u>P</u>; residues 26 to 31). Similarly to SseF, expression of SseG single or triple alanine mutations in the conserved residues of the putative CBD motif (SseG_P26A_, SseG_L28A,_ SseG_P31A_ and SseG_P26L28P31A_) led to a decrease in SseG secretion. Secretion of SseG_L28A_ and SseG_P26L28P31A_ significantly decreased to 9 ± 4 % and 7 ± 2 %, respectively, when compared with wild-type SseG (Fig. 5F). All SseG mutant proteins had similar expression levels (Fig. 5F).

In addition to the analysis of the SseF and SseG alanine mutants in *Salmonella*, we measured the melting temperature (T_m_) of purified SscB (Fig. 3B), together with peptides containing mutated CBD motifs of SseF and SseG (_pep_SseF_P32I34P37A_, _pep_SseG_P26L28AP31A_). We observed that the mutated peptides had a lower T_m_ than their non-mutated version and that of SscB alone (SscB, T_m_ = 38.62 °C; _pep_SseF, T_m_ = 40.79 °C; _pep_SseF_P32I34P37A_, T_m_ = 34.89 °C; _pep_SseG, T_m_ = 40.38 °C; and _pep_SseG_P26L28AP31A_, T_m_ = 37.83 °C). In summary, we identified PXL/IXXP motifs in both SseF (<u>P</u>EIPA<u>P</u>; residues 32 to 37; Fig. 5C) and SseG (<u>P</u>S<u>L</u>PH<u>P</u>; residues 26 to 31; Fig. 5G), respectively, and found that these residues are key for protein-protein interactions that stabilize SscB and allow for efficient secretion of both SseF and SseG.

### Structural conservation of class II chaperones is not synonymous with secretion hierarchy

The striking conservation of the structure and mode of interaction of both SscB/SseF and SscB/SseG with class II chaperone/translocators complexes (Fig. 4D and 4E) led us to investigate the conservation of chaperone functions. Using Alphafold2, we compared the predicted structures of SscB with those of SicA and SscA, the class II chaperones of the translocators of the *Salmonella* T3SS-1 and T3SS-2, respectively. The structures of the three proteins had high confidence scores (pLDDT > 95) (Fig. S7) and revealed typical α-helical structures of class II chaperones and a predicted pocket for the binding of their respective substrates (Fig. 6A). The predicted substrate-binding regions of SscB and SicA are composed of only aromatic hydrophobic amino acids (tyrosine and tryptophan) that coordinate the interaction with a chaperone-binding motif. Chaperones of other bacteria were shown to have a similar amino acid composition (Fig. 4D). In contrast, in SscA this region is more chemically diverse comprising tyrosine, threonine and a methionine. (Fig. 6A). Based on their structural similarity to SscB, we wondered whether SscA and SicA could complement some of its functions. Therefore, we analysed *Salmonella sscB* mutant strains co-expressing SseF paired with SscA and SicA, respectively. SicP, a class I chaperone of the effector SptP was used as a negative control. The co-expression of SicA with SseF resulted in low SseF secretion (11 ± 3 %), while no secretion signal was detected when SseF was co-expressed with SscA or SicP (Fig. 6B). However, the signal detected for SseF co-expressed with SicA was independent of the T3SS-2, as it was also detected in T3S-deficient strains (*sscB ssaV*). To rule out secretion via T3SS-1, we performed a split-NanoLuc secretion assay in SPI-1 inducing conditions, using as a positive control the T3SS-1 effector SipA. Comparison of the T3S-proficient (*sscB*) and T3S-deficient (*invA*) strains for T3SS-1 revealed no differences, indicating that low SseF secretion when co-expressed with SicA is both T3SS-1 and T3SS-2 independent (Fig. 6C). In parallel, we analysed whether a class II chaperone of translocators could alter the secretion hierarchy and change SseF from a late to an intermediate substrate. We used the deletion mutant of one of the components of the T3SS-2 gatekeeper complex, SsaL, which is described to oversecrete effectors (late substrates) and abolish secretion of translocators (intermediate substrates) (Yu et al., 2010) (Yu et al., 2018). If SseF was secreted as an intermediate substrate in the presence of SicA, we should observe a reduction in its secretion. This was not the case: while SseF, co-expressed with SscB, was oversecreted in the *ssaL* mutant, no reduction was observed when co-expressed with SicA (Fig. 6B). Additionally, we also observed that SseG co-expressed with SscB, behaved as a bona fide effector and was oversecreted in the *ssaL* mutant (Fig. S8). Lastly, since a low secretion signal was observed for SseF when paired with SicA in all the tested mutants (*sscB*, *sscB ssaV*, *sscB ssaL*) (Fig. 6 B), we assessed whether SicA could be stabilising SseF. In the presence of SicA, the stability of SseF was similar to that of SseF alone (Fig. 2C and S4). Overall, SscA and SicA could not facilitate the secretion of SseF, thus highlighting that each chaperone is highly specialised to its own substrates, despite structural conservation.

**Figure 6:**
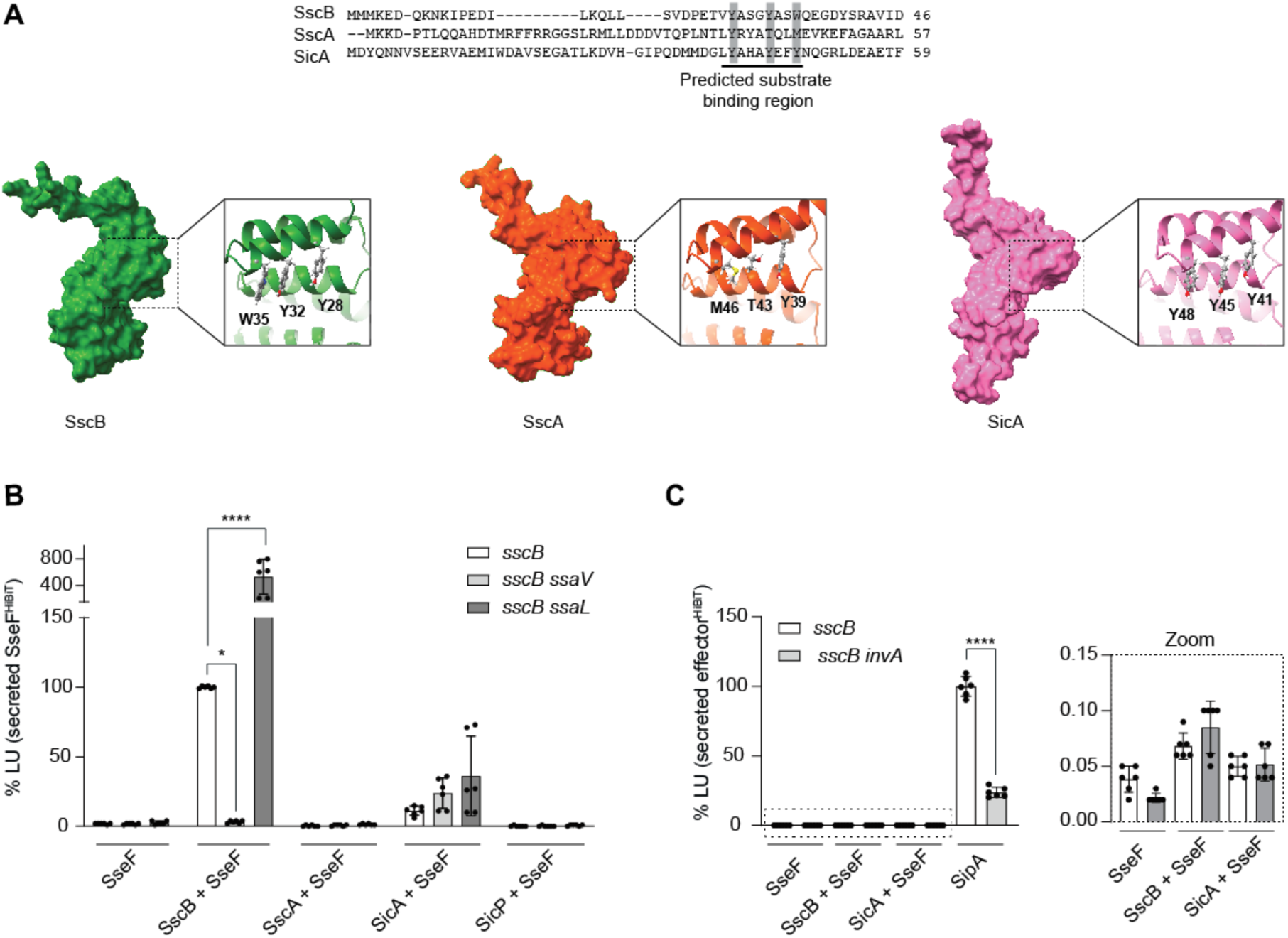
Conservation of the function of class II chaperones. **(A)** Alignment of the sequences of SscB and the class II chaperones of *Salmonella* SscA (T3SS-2) and SicA (T3SS-1) containing the predicted substrate**-**binding region. Highlighted, in grey, are the residues predicted to coordinate the interaction with the CBD motif of the substrate. Structure predictions were obtained using Alphafold2 Multimer with a detail view of the substrate**-**binding pocket. **(B)** T3S-proficient (*sscB*), T3S-deficient (*sscB ssaV*) and gatekeeper mutant (*sscB ssaL*) *Salmonella* strains, harbouring plasmids co-expressing chaperone and effectors, under rhamnose-inducible promoter, were grown under SPI-2-inducing and secreting conditions. Bacterial cultures were collected, and secretion was analysed using the split-NanoLuc T3S assay. The secretion of plasmid-derived SseF^HiBiT^ was analysed by luminometry. The luminescence signal was compared to the T3S-proficient strain (*sscB*) expressing SscB and SseF (SscB + SseF). **(C)** *Salmonella* strains proficient (*sscB*) and deficient (*sscB invA*) in secretion via T3SS-1 were grown under SPI-1-inducing and secreting conditions. Bacterial cultures were collected, and secretion was analysed using the split-NanoLuc T3S assay. The secretion of plasmid-derived effector^HiBiT^ was analysed by luminometry. The luminescence signal was compared to the T3S-proficient strain (*sscB*) expressing SipA. Statistical significance is indicated as follows: * P<0.05, ****P<0.0001, further description of the statistical analysis can be found in Supp. Table 3

## Discussion

The targeting of proteins to T3SSs is a process critical for the ability of pathogenic bacteria to cause infection. Protein secretion relies on specific T3S signals that facilitate the delivery of bacterial proteins into host cells. Secretion of transmembrane T3S substrates poses an additional challenge as their hydrophobic transmembrane segments are prone to aggregation, degradation and mistargeting to bacterial inner membrane. T3S chaperones solve this problem by binding and masking the transmembrane segments of the substrates. Nevertheless, to ensure efficient secretion, the steps preceding secretion must occur rapidly and be highly regulated. Our study provides further insights into the sequence of events that culminate in the secretion of the *Salmonella* transmembrane effectors, SseF and SseG, assisted by their chaperone SscB. We show that from gene location to the formation of a chaperone/effector complex, to protein stabilisation and secretion, a series of interconnected and interdependent events must occur. We also uncovered that the SscB/SseF and SscB/SseG chaperone-effector pairs structurally resemble class II chaperone-intermediate substrate pairs while the functionally behave like class I chaperone-effector pairs. This is the first description of a class II chaperone that binds effectors, thus challenging the paradigm that the hierarchy of secretion correlates with chaperone structure.

The genes encoding transmembrane effectors SseF and SseG and their chaperone SscB are located adjacently in the operon *sscB-sseF-sseG*. We observed that changing gene order from *sscB-sseF* to *sseF-sscB* significantly decreased the secretion and injection of SseF and the ability of *Salmonella* to cause infection. These findings align with previous observations for other T3S chaperone/effector pairs. For instance, in *Escherichia coli,* gene location of the effector/chaperone pair *tir-cesT* is essential for maintaining mRNA stability (Elbaz et al., 2019), while in *Salmonella*, gene location of SPI-1 chaperone/effector pair SicP/SptP is required for coupled translation (Button and Galán, 2011). However, for SscB/SseF a different regulatory mechanism may be at play. It was previously reported that SscB and SseF only interact when co-expressed from a bicistronic vector, as no interaction was detected when each protein was expressed independently in separate bacterial strains after their bacterial lysates were combined (Dai and Zhou, 2004). Our results obtained from expression inside the same *Salmonella* suggest otherwise. Expression of SscB in *trans* stabilized SseF, indicating formation of SscB/SseF complexes and resulting in oversecretion of SseF. SseF oversecretion was not, however, the result of a positive regulatory effect of SscB as previously reported for other T3SS chaperones (Darwin and Miller, 2000; Fei et al., 2021; Francis et al., 2001). Collectively, our results suggest that SscB facilitates SseF secretion by co-translational interaction, thereby preventing co-translational mistargeting by the SRP. However, binding of neither of these two competing targeting factors seems to be particularly faithful as a surplus of in-trans expressed SscB results in its strongly increased type III secretion.

Our results further show that the structural stability of SscB, SseF and SseG is another factor controlling secretion. SscB has a melting temperature (T_m_ = 38.26°C) close to the optimal growth temperature of *Salmonella* and the body temperature of many of their hosts. This T_m_ is low compared to the T_m_ other T3S chaperones (*E. coli* CesT, 63 °C; *Aeromonas* AcrH, 52 °C; *Pseudomonas* PcrH, 48 °C;) (Ramu et al., 2013; Schoehn et al., 2003; Tan et al., 2009). Addition of a peptide containing the chaperone-binding domain (CBD) of SseF or SseG was sufficient to increase the melting temperature of SscB. The melting temperature as proxy for stability indicates that SscB alone is not stable under physiological conditions. This was confirmed in *Salmonella*, where SscB was not stable by itself, and was only stabilized by co-expression with full-length SseF but not with SseG. Furthermore, SseF was unstable in the absence of SscB, as previously observed (Dai and Zhou, 2004), whereas SseG was stable by itself. Similar stability relationships have been described for translocators and their cognate chaperones. In *Shigella,* chaperone IpgC is necessary for stable expression of translocator IpaB but not IpaC (Ménard et al., 1994), and in *Yersinia,* chaperone SycD is required for YopB but not YopD stability (Neyt and Cornelis, 1999). Furthermore, the *sscB-sseF-sseG* operon organization recapitulates the *chaperone-translocator-translocator* genomic organization, and in each case the chaperone is more critical for stabilising the substrate encoded in closest genomic proximity. Interestingly, in all cases, the more hydrophobic substrate is encoded in proximity to the chaperone (SseF, IpaB and YopB) and the less hydrophobic encoded farther downstream (SseG, IpaC, YopD). This conservation suggests that within class II chaperone operons, genomic proximity correlates with the degree of chaperone co-dependence for stability.

SscB has long been annotated as a “CesD/SycD/LcrH family type III secretion system chaperone”, belonging to the class II chaperones that bind translocators (Cirillo et al., 1998), yet SseF and SseG are bona fide effectors. To our knowledge, this inconsistency has never been paid attention to. We report that SscB contains only α-helices, supporting its similarity to known class II chaperones and not to class I chaperones, which typically exhibit an α/β sandwich fold. Furthermore, purified SscB formed dimers, while the SscB/SseF complex exhibited a 1:1 stoichiometry, suggesting dissociation of the dimer upon interaction with SseF. This behaviour was observed for other class II T3S chaperones; for *Pseudomonas aeruginosa* class II chaperone PcrH, its dimerization was suggested to be an intermediate state prior to the formation of the chaperone/translocator complex with translocators PopB and PopD (Tomalka et al., 2013). To further characterize the SscB/SseF and SscB/SseG complexes, we mapped their interaction interfaces. The N-terminus of SscB was found to interact with both SseF and SseG. Although the N-terminus of a class II chaperone is not required for its stability or dimerization (Job et al., 2010; Singh et al., 2013), it plays an essential role for chaperone/translocator interaction, as the deletion of the N-terminus of *Aeromonas* chaperone AcrH prevents formation of the chaperone/translocator (AcrH-AopB) complex (Nguyen et al., 2015). Another conserved feature of class II chaperones is a substrate-binding pocket that engages with a conserved CBD motif within the cognate substrate. We identified such CBD motifs in SseF (PEIPAP, residues 32 - 37) and SseG (PSLPHP, residues 26 - 31). Alanine substitution of the three highly conserved residues of the motif disrupted the interaction with SseF and halted its secretion; the same was observed for SseG. Furthermore, mutagenesis of the second conserved residue in SseF (isoleucine, I34) and SseG (leucine, L28) was sufficient to completely abolish the secretion of both proteins. These results are consistent with the observation that equivalent mutations in *Pseudomonas aeruginosa* translocator PopB reduced the binding affinity for its chaperone PcrH (Frankling et al., 2023). Beyond the CBD, the first and also the second transmembrane segments of SseF interacted with SscB. No interaction with SscB was detected across multiple positions within SseG transmembrane segments. Taken together, our data show that the SscB/SseF and SscB/SseG complexes share structural features and mode of interaction characteristic of class II chaperone/translocator complexes, further reinforcing that SscB, SseF and SseG behave as class II chaperones and translocators prior to secretion.

Class II chaperones have been described to interact exclusively with translocators, and class I chaperones only with effectors, giving rise to the belief that chaperone class is a determinant for both substrates targeting to the injectisome and hierarchy of secretion. This view was supported by the structural conservation within each chaperone class, suggesting that the mode of interaction of the chaperone with the injectisome components was conserved and could control the timing of secretion. SscB directly challenges this hypothesis as it has the hallmark features of a class II chaperone yet delivers effectors as late substrates and not translocators as intermediate ones.

The striking structural conservation between SscB/SseF and SscB/SseG complexes and canonical translocator complexes led us to investigate whether structurally similar chaperones could be interchangeable. SscB and the *Salmonella* class II chaperones SicA (SPI-1) and SscA (SPI-2) were structurally similar, with SscB and SicA binding regions being composed exclusively of aromatic hydrophobic residues, as other bacterial class II chaperones. However, neither SicA nor SscA could stabilise SseF or restore its secretion, nor could either chaperone alter the secretion hierarchy of SseF from a late to an intermediate substrate. Thus, despite structural similarities between the chaperones, substrate recognition is highly specific both within the same bacterium and across different T3SSs, as previously demonstrated by the inability of *Pseudomonas* translocators to substitute for their *Yersinia* counterparts (Armentrout and Rietsch, 2016).

SscB, SseF, and SseG recapitulate the structure and behaviour of class II chaperones and their translocators until the moment of secretion, prompting the question on the biological relevance of this structural conservation. We hypothesize that these similarities reflect the shared function to bind transmembrane substrates encoded in genomic proximity, thereby preventing premature interaction of transmembrane proteins that, upon secretion into host cells, interact with each other. This applies to both translocators and the transmembrane effectors SseF and SseG. The finding that a class II chaperone delivers effectors rather than translocators raises another question regarding how bacteria discriminate between translocators and effectors and, therefore, the hierarchy of secretion. Our works suggests that the structural features of class II chaperones are not sufficient for such discrimination, contrary to the general observation that hierarchy of secretion correlates with shared chaperone structures (Parsot et al., 2003). Therefore, other determinants must be involved. One model proposes a role for N-terminal secretion signals, however, it was demonstrated that secretion signals can also be encoded at the C-terminus of the substrate or near the CBD (Tomalka et al., 2012). Another model attributes discrimination to differences in affinities for the gatekeeper (Portaliou et al., 2017) or for other injectisome components, such as ATPase or the export apparatus. Additional layers of specificity must therefore reside within the substrate, the chaperone–substrate interface, or their interactions with injectisome components. Further studies will be necessary to fully elucidate the function and specificity of class II chaperones in a broader mechanistic context.

## Material and Methods

### Bacteria and cell lines

*Escherichia coli* NEB 10β (New England Biolabs) was used for construction of plasmids. *E. coli* LEMO21 (DE3) (Wagner et al., 2008) and *E. coli* BL21 (DE3) were used for protein purification. *E. coli* strains were routinely grown at 37 °C in liquid or solid lysogeny broth (LB) medium with the appropriate antibiotics and supplements. All *Salmonella* enterica Typhimurium strains were derived from the NCT12023 strain. Bacterial cultures were supplemented as required with kanamycin (50 μg/mL). *Salmonella* mutant strains were constructed by allelic exchange using the suicide plasmids listed in Table S4. HeLa cells constitutively expressing LgBiT (Hela-LgBiT) (Westerhausen et al., 2020) were maintained in high-glucose Dulbecco’s modified Eagle Medium (DMEM) supplemented with heat-inactivated 10% (v/v) fetal bovine serum (FBS) at 37 °C in a humidified atmosphere of 5% (v/v) CO_2_. Cells were routinely checked for *Mycoplasma* by PCR.

### Plasmids

The plasmids and primers used in this work and their main characteristics and construction details are described in Table S4. Molecular cloning was performed by standard Gibson cloning, and site-directed mutagenesis was performed following the Quick-Change protocol (Stratagene) using KOD polymerase (Novagen). The accuracy of the nucleotide sequence of all the plasmids was confirmed by DNA sequencing (Eurofins).

### Protein purification of SscB

Recombinant protein 6His-TEV-SscB was expressed in *E. coli* LEMO21 (DE3) (Wagner et al., 2008). Bacteria were grown overnight in Luria-Bertani broth (LB) supplemented with kanamycin (50 µg/mL) and chloramphenicol (20 µg/mL). Overnight cultures were diluted to an OD_600_ of 0.05 in Terrific broth (TB) and incubated in 1 L shaker flasks at 37°C, 180 rpm. When OD_600_ reached 0.4, isopropyl-D-thiogalactopyranoside (IPTG) was added to a final concentration of 0.4 mM. Cultures were incubated at 18 °C, 180 rpm for 16 h. Bacteria were harvested, and pellets were resuspended in lysis buffer (50 mM Tris-HCl, 200 mM NaCl, 5 % glycerol, pH 8) supplemented with DNase I (10 µg/mL), lysozyme (10 µg/mL), protease inhibitor cocktail (1x; Sigma Aldrich) and EDTA (1mM). Cells were lysed using the French Press (2 times, 10 000 psi) and 1 mM MgCl_2_ was added. The lysate was centrifuged at 24 0000 *x g*, for 20 min at 4 °C. Supernatant was collected and ultracentrifuged at 45 000 *x g*, for 45 min at 4 °C. The resulting supernatant was loaded in a pre-equilibrated HisTrap HP column (Cytiva) and further washed with washing buffer (50 mM Tris-HCl, 200 mM NaCl, 5 % glycerol, 10 mM imidazole, pH 8). Protein was eluted in an imidazole gradient elution buffer (50 mM Tris-HCl, 200 mM NaCl, 5 % glycerol, 250 mM imidazole, pH 8). The eluted protein was digested with TEV protease and dialysed against the dialysis buffer (50 mM Tris-HCl, 200 mM NaCl, 5 % glycerol, pH 8). The dialysed eluate was loaded on a HisTrap HP column (Cytiva) pre-equilibrated with lysis buffer: The unbound fraction (untagged SscB) was collected, and the protein was purified using size exclusion chromatography column (SEC) Hiload 16/600 Superdex 200 pg (Cytiva) using lysis buffer. Purified protein was concentrated using Amicon Centrifugal Filter cut-off 3 kDa, and stored at −80 °C.

### Protein purification of SscB/SseF

The complex 6His-TEV-SscB/SseF-TEV-StrepII was expressed in *E. coli* BL21 (DE3). Bacteria were grown overnight in Luria-Bertani broth (LB) supplemented with kanamycin (50 µg/mL). Overnight cultures were diluted to an OD_600_ of 0.05 in Terrific broth (TB) and incubated in 1 L shaker flasks at 37°C, 180 rpm. When OD_600_ reached 0.4, isopropyl-D-thiogalactopyranoside (IPTG) was added to a final concentration of 0.4 mM. Cultures were incubated at 18 °C, 180 rpm for 16 h. Bacteria were harvested, and pellets were resuspended in lysis buffer (50 mM Tris-HCl, 200 mM NaCl, 5 % glycerol, pH 8) supplemented with DNase I (10 µg/mL), lysozyme (10 µg/mL), protease inhibitor cocktail (1x; Sigma Aldrich) and EDTA (1mM). Cells were lysed using the French Press (2 times, 10 000 psi) and 1 mM MgCl_2_ was added. The lysate was centrifuged at 24 0000 *x g*, for 20 min at 4 °C. Supernatant was collected and ultracentrifuged at 45 000 *x g*, for 45 min at 4 °C. The resulting supernatant was loaded on a pre-equilibrated StrepTrap HP column (Cytiva) and washed using lysis buffer. The protein complex was eluted using (50 mM Tris-HCl, 200 mM NaCl, 5 % glycerol, 2.5 mM desthiobiotin, pH 8). The protein was purified using size exclusion chromatography column (SEC) Hiload 16/600 Superdex 200 pg (Cytiva) with lysis buffer. Purified protein was concentrated using Amicon Centrifugal Filter cut-off 3 kDa, and stored at −80 °C.

### Size-exclusion chromatography with multi-angle light scattering (SEC-MALS)

SEC-MALS experiments were performed using a Superdex 75 Increase 10/300 Gl column (Cytiva) coupled to a miniDAWN Tristar Laser photometer (Wyatt) and a RI-2031 differential refractometer (JASCO). Fifty microliters of protein samples were loaded to the SEC column, equilibrated with SEC buffer (50 mM Tris-HCl, 200 mM NaCl, 5 % glycerol, pH 8), and separated using a flow rate of 0.5 ml min−1. Data analysis was carried out with ASTRA v7.3.0.18 software (Wyatt).

### Nano differential scanning fluorimetry (Nano-DSF)

Protein samples were heated from 20 to 80 °C, with a temperature gradient of 0.4 °C min−1. Melting temperatures were calculated from changes in the fluorescence ratio (350/330 nm), using PR.Stability Analysis v1.0.3 software (NanoTemper Technologies GmbH). To assess peptide-induced effects on SscB stability, peptides were diluted to final assay concentration of 100 μM into 0.2 mg/mL SscB (in lysis buffer). Samples were incubated for 10 min at room temperature before loading of standard capillaries (NanoTemper Technologies GmbH)

### Circular Dichroism (CD)

CD spectra were recorded on a Jasco J-810 spectrometer, with spectral scan window of 200-250 nm, with a data pitch of 0.1 nm and a scan speed of 100 nm min-1. CD spectra are an average of 5 individual scans. Melting curves were measured from 20 to 100 °C, recording the ellipticity at λ = 222 nm every 0.5 °C, with a temperature gradient of 1 °C/min.

### *In vivo* photocrosslinking

*Salmonella* Typhimurium strains were transformed with pT10-derived and pSUP plasmids (Table S4). Bacteria were grown overnight in LB broth supplemented with kanamycin (50 µg/mL) and chloramphenicol (20 µg/mL). Bacteria were diluted to an OD_600_ of 0.1 in SPI-2 inducing and secreting medium (PCN, pH 5.8) supplemented with 1 mM rhamnose and with the artificial amino acid para-benzoyl phenylalanine (pBpa) to a final concentration of 1 mM. Cultures were grown at 37 °C for 6 h, 180 rpm. 8 ODU of bacterial cells were harvested and washed once with ice-cold PBS. Bacteria were resuspended in 2 mL of PBS, and 1 mL (4 ODU) was kept on ice (-UV). The other 1 mL (4 ODU) was transferred to a well in a 6-well plate and irradiated with λ = 365 nm UV on a UV transilluminator table for 30 min (+UV). Bacteria were pelleted and resuspended in 1 x SB buffer and boiled at 95 °C, 10 min. Samples were analysed by SDS-PAGE and immunoblot.

### *In vivo* protein stability assay

Overnight cultures of *Salmonella* Typhimurium were diluted to an OD_600_ of 0.2 in PCN medium (pH 5.8), and incubated at 37 °C, 180 rpm. When OD_600_ reached 0.5, rhamnose was added to the final concentration of 1mM. After 3 h of induction, protein synthesis was halted by adding 100 µg/mL of chloramphenicol. Samples were taken every 15 min. Bacteria were harvested and resuspended in 1 x SB buffer and boiled at 95 °C, 10 min. Samples were analysed by SDS-PAGE and immunoblot.

### Split-NanoLuc based T3S Assay under SPI-2 inducing conditions

Overnight cultures of *Salmonella* Typhimurium were diluted to an OD_600_ of 0.1 in PCN medium (pH 5.8), and if necessary supplemented with 1 mM rhamnose, and incubated at 37 °C, 180 rpm for 6 h. Then, 25 µL of the culture was transferred to a 384-well plate, and to each well was added 25 µL of reaction solution containing extracellular buffer, LgBiT (1:100), and extracellular substrate (1:50) (Promega). The luminescence was measured using a Tecan Spark reader.

### Split-NanoLuc based T3S Assay under SPI-1 inducing conditions

Overnight cultures of *Salmonella* Typhimurium were diluted to an OD_600_ of 0.1 LB, 0.3 M NaCl, supplemented with 1 mM rhamnose and incubated at 37 °C, 180 rpm for 5 h. Then, 25 µL of the bacterial culture was transferred to a 384-well plate. To each well, 25 µL of reaction solution containing extracellular buffer, LgBiT (1:100), and extracellular substrate (1:50) (Promega) were added. The luminescence was measured using a Tecan Spark reader.

### Split-NanoLuc based injection assay

HeLa-LgBiT cells were seeded in white 96-well plates with glass bottom (1 x 10^4^ cells/well). The day after seeding, overnight cultures of *Salmonella* Typhimurium were diluted to an OD_600_ of 0.1 and grown until OD_600_ of 0.9. Bacterial inoculum was prepared in Hanks’ Balanced Salt Solution (HBSS), added to HeLa-LgBiT cells at a multiplicity of infection (MOI) of 100 and incubated for 1 h at 37 °C in a humidified atmosphere of 5 % (v/v) CO_2_. The inoculum was removed and replaced by DMEM supplemented with 10 % (v/v) FBS and 100 μg/mL of gentamicin for 1 h and then with 16 μg/mL of gentamicin. After 16 h of infection, the cells were washed and 100 µL of HBSS with DrkBiT (1:1 000) was added. Next, 25 µL of reaction buffer containing NanoGlo Live Cell buffer and furimazine substrate (1:20) were added to the wells. Luminescence was measured using a Tecan Spark reader.

## Acknowledgements

This work was funded by infrastructural measures of the cluster of excellence EXC2124 Controlling Microbes to Fight Infections (CMFI). P.F. and S. Schroth. were supported by the DZIF MD Program TI 07.003. We acknowledge support from the Open Access Publication Fund of the University of Tübingen.

We thank Michael Hensel for providing the published *Salmonella Typhimurium* NCTC 12023 strain, Nelli Deobald for assistance with SscB/SseF purification, Andrea Eipper for technical support in the laboratory, and Libera Lo Presti for critically reading the manuscript.

## Author contributions

S.V.P. and S.W. conceived the experiments. S.V.P., P.F., S. Schroth, J.J., E.P., S. Schminke conducted the experiments. S.V.P., P.F., S. Schroth., J.J., Schminke, M.H. and S.W. analysed the results. S.V.P. and S.W. wrote the manuscript, which all authors revised.

## Supplementary material (Text/Figures)

### Supplementary Text 1: Optimisation of split-NanoLuc-based for detecting the secretion via T3SS-2 in bacterial cultures

We successfully detected SseF^HiBiT^ secretion in the culture supernatant (SN) using classic TCA precipitation method and the split-NanoLuc T3S assay (Fig. S2A and S2B). In the T3S-deficient strains (*ssaV*), secretion was not detected. In addition, we observed that SseF expression under its endogenous promoter (P_sseA_) resulted in higher protein levels and secretion compared to expression under the rhamnose-inducible promoter (P_rha_) (Fig. S2A and S2B). Most importantly, in bacterial cultures, a marked increase in the secretion signal was detected, consistent with previous observations showing that after secretion, SseF is not only found in the supernatant but also attached to the bacterial surface (Hansen-Wester et al., 2002). Even in this case, we detected higher secretion when expressing the proteins from an endogenous promoter (Fig S2B). Nevertheless, the signal-to-noise ratio (S/N) was similar for both promoters when using bacterial cultures (Fig S2C). These results demonstrate that the split-NanoLuc T3S assay can effectively detect SseF secretion through the T3SS-2 of *Salmonella,* especially from whole bacterial cultures.

---

**Figure S1:**
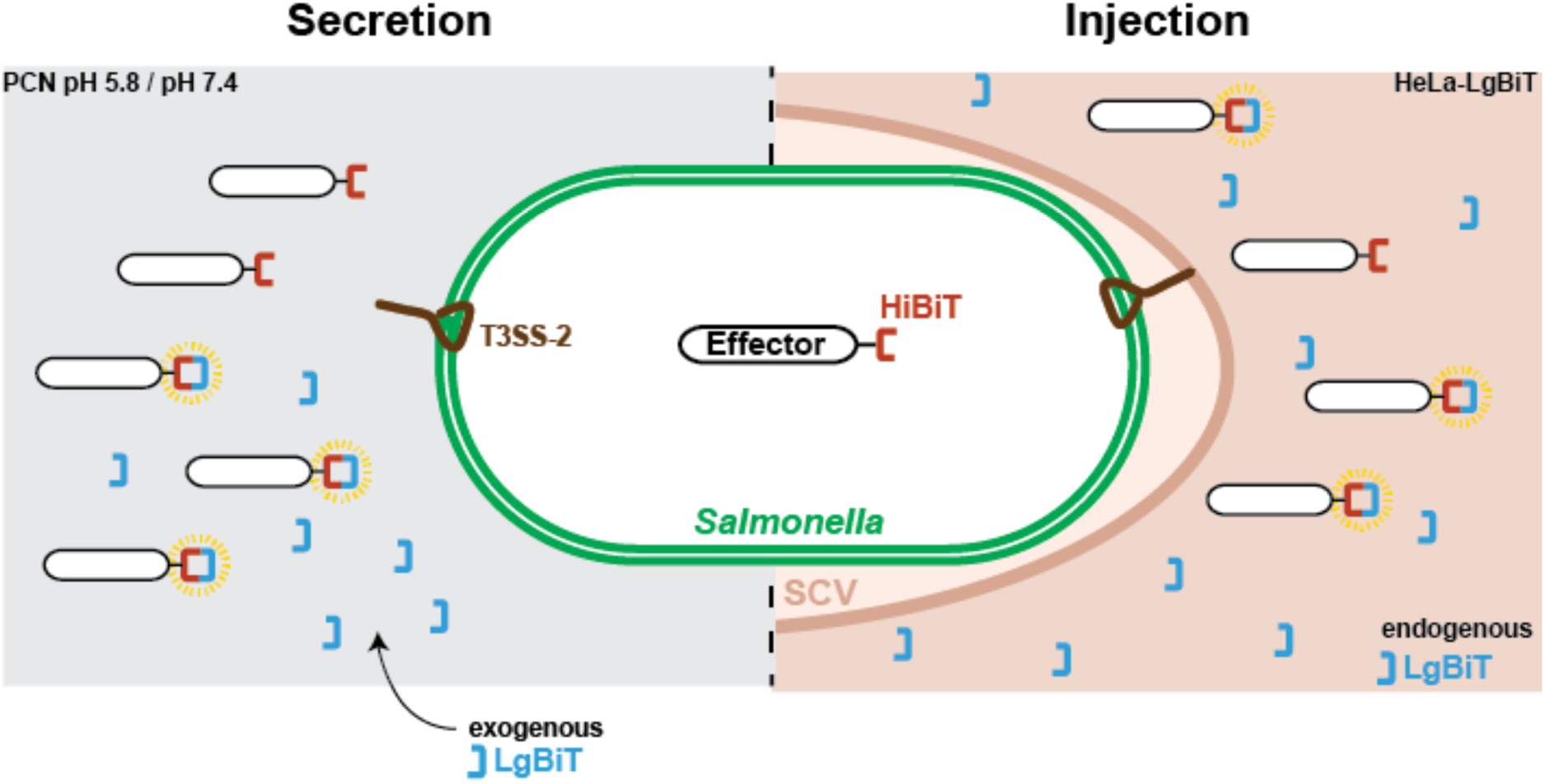
Split-NanoLuc-based assay for detecting the secretion and injection of effectors via the T3SS-2 of *Salmonella*. To assess type III secretion (left), *Salmonella* strains expressing effector^HiBiT^ are grown in SPI-2 inducing and secreting media (PCN, pH 5.8) for 6 h at 37°C. Next, bacterial cultures or bacterial supernatant are collected, and LgBiT and NanoLuc substrate are added. If effector-HiBiT is secreted, the reconstituted HiBiT/LgBiT complex catalyses a reaction that produces a luminescence signal. To assess the injection of effectors into host cells (right), HeLa cells constitutively expressing LgBiT (HeLa-LgBiT) are infected with *Salmonella* expressing effector^HiBiT^. After infection, the substrate and DrkBiT are added. If the effector-HiBiT is translocated, the complementation of HiBiT/LgBiT leads to a luminescence signal.

**Figure S2:**
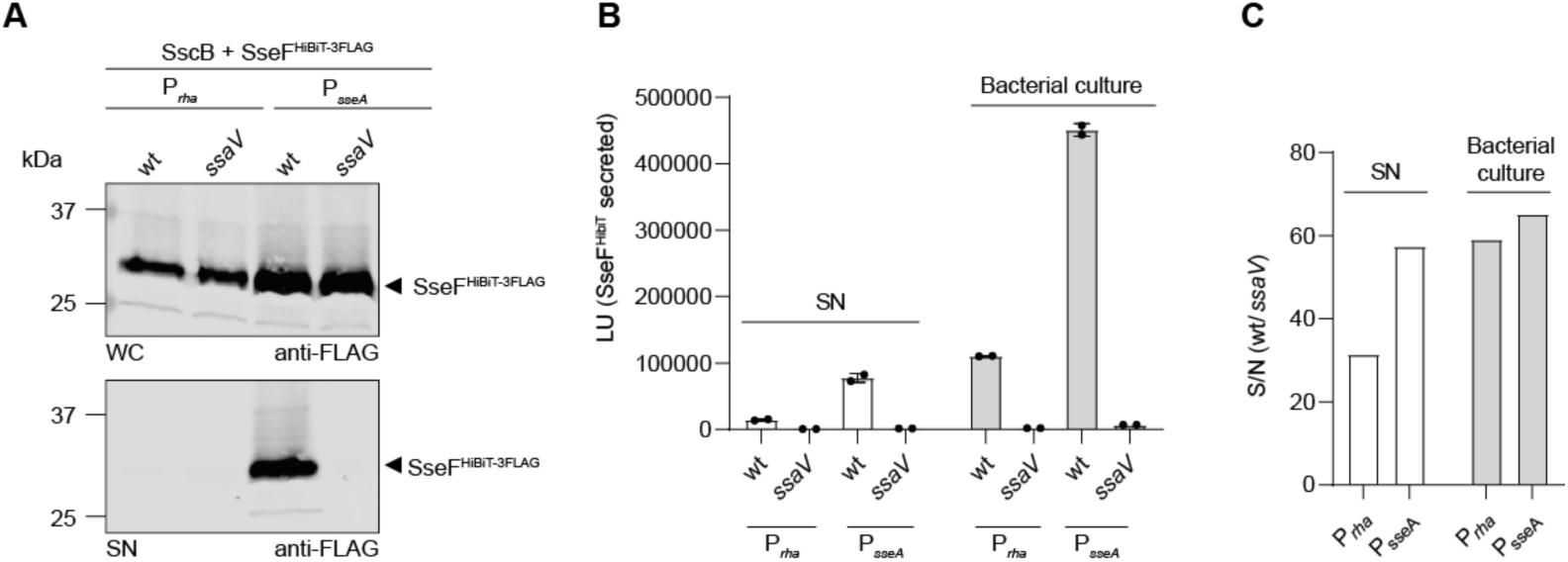
Optimisation of split-NanoLuc-based for detecting secretion via T3SS-2. **(A)**, **(B)** and **(C)** *Salmonella* T3S-proficient (wt) and T3S-deficient (*ssaV*) strains expressing SseF^HiBiT^ from endogenous (P_sseA_) or rhamnose-inducible (P_rha_) promoter were grown in SPI-2 inducing and secreting media (PCN, pH 5.8) for 6 h at 37°C. **(A)** Whole-cell extracts (WC; non-secreted proteins) and culture supernatant (SN; secreted proteins) were collected. The proteins in the culture supernatant were precipitated using TCA. Proteins in whole-cell extracts and culture supernatant were analysed by SDS-PAGE and immunoblot using the indicated antibodies. **(B)** LgBiT and NanoLuc substrate were added to culture supernatant (SN) or bacterial cultures (bacteria + supernatant). Secreted SseF^HiBiT^ was measured by luminometry. **(C)** Signal-to-noise ratio (wt/*ssaV*) for luminescence signal of secreted SseF^HiBiT^ in bacterial supernatant (SN) and bacterial cultures.

**Figure S3.**
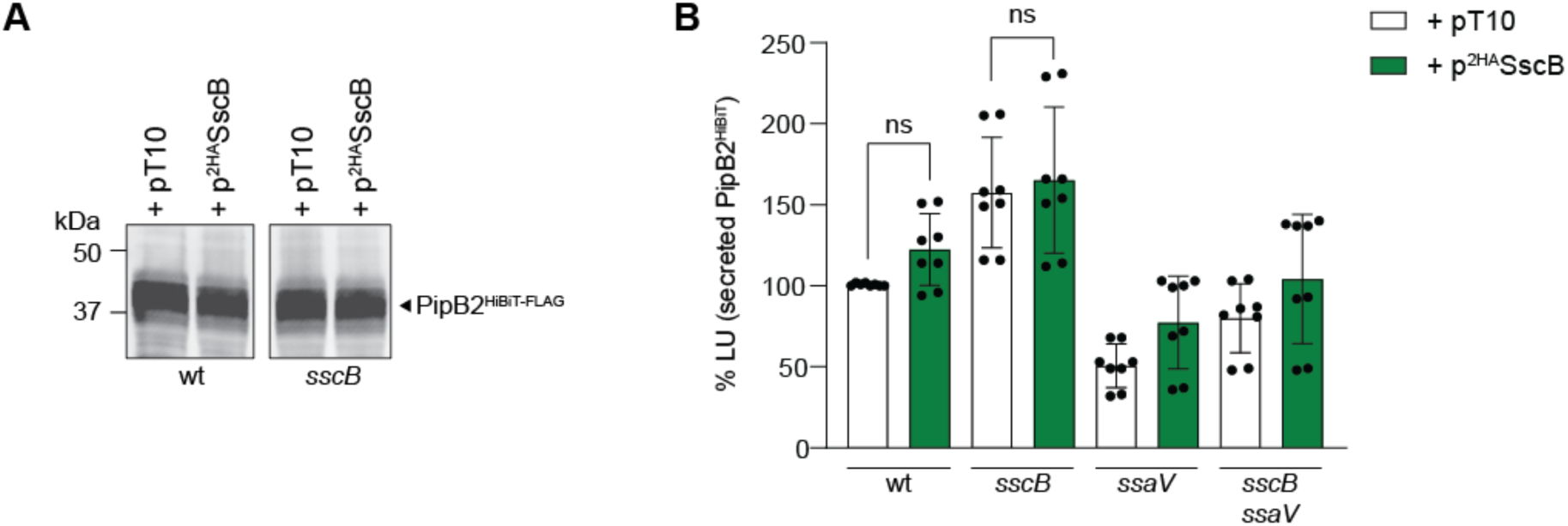
**Effect of SscB on the secretion of effector PipB2**. *Salmonella* wild-type (wt) and *sscB* mutant strains expressing PipB2^HiBiT-3FLAG^ from the chromosome were complemented with empty pT10 or p^2HA^SscB. Bacteria were grown under SPI-2-inducing and secreting conditions for 6 h. **(A)** Bacteria were harvested and analysed by SDS-PAGE and immunoblot using anti-FLAG antibody. **(B)** Bacterial cultures were collected, and secretion was analysed using the split-NanoLuc secretion assay. The secretion of chromosome-derived PipB2^HiBiT^ was analysed by luminometry. The luminescence signal was compared to the wild-type strain harbouring the pT10 plasmid. Data are the mean ± SD from 4 independent experiments. P-values were obtained by one-way ANOVA analyses; ns (not significant) P > 0.05.

**Figure S4:**
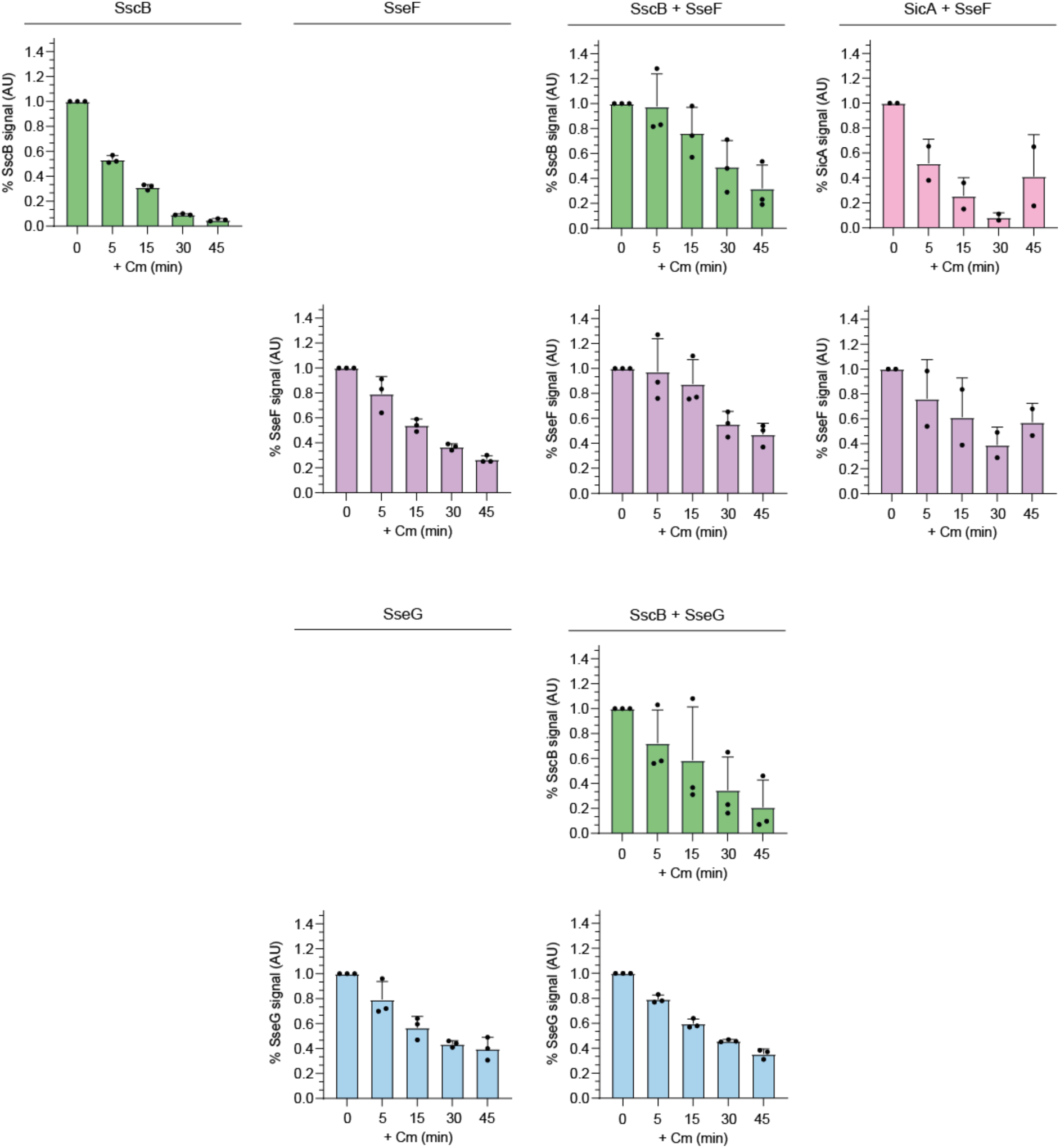
Quantification of protein stability. Protein stability was analysed in T3S-deficient (*sscB ssaV*) *Salmonella* strains expressing chaperone/effector^HiBiT^ pairs from a rhamnose-inducible (P_rha_) promoter. Bacteria were grown under SPI-2-inducing conditions and protein expression was induced when OD_600_ reached 0.5. After 3h of induction, chloramphenicol (Cm) was added and samples were harvested every 15 min. Whole-cells extracts were analysed by SDS-PAGE immunoblot using antibodies against HA and FLAG (see Fig. 2C). Quantification was performed using Image Studio Lite and normalised to a control band. Further, the normalised signal was compared to time zero. Time zero (0 min) corresponds to time before the addition of chloramphenicol. Data are the mean ± SD from 3 independent experiments, except for SicA + SseF (n =2)

**Figure S5:**
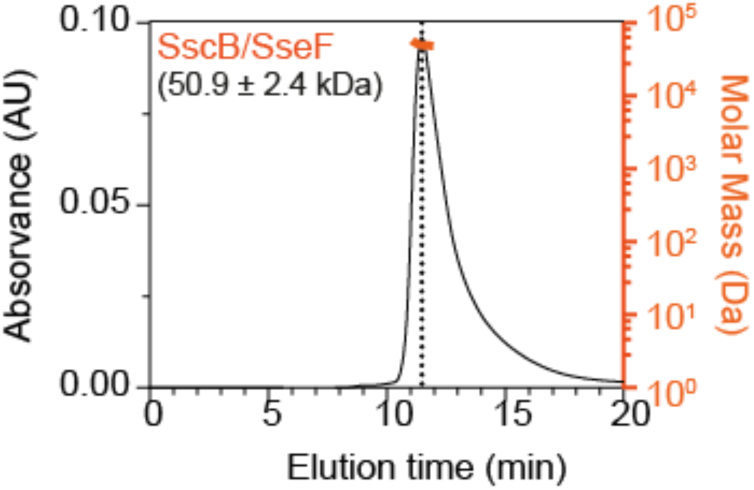
SEC-MALS chromatogram of purified tagged-SscB/SseF (^6His^SscB/SseF^StrepII^) complex. The predicted molecular mass of ^6His^SscB is 19.6 kDa and of SseF^StrepII^ is 29.2 kDa. The peak corresponds to the SscB/SseF complex in a 1:1 stoichiometry.

**Figure S6:**
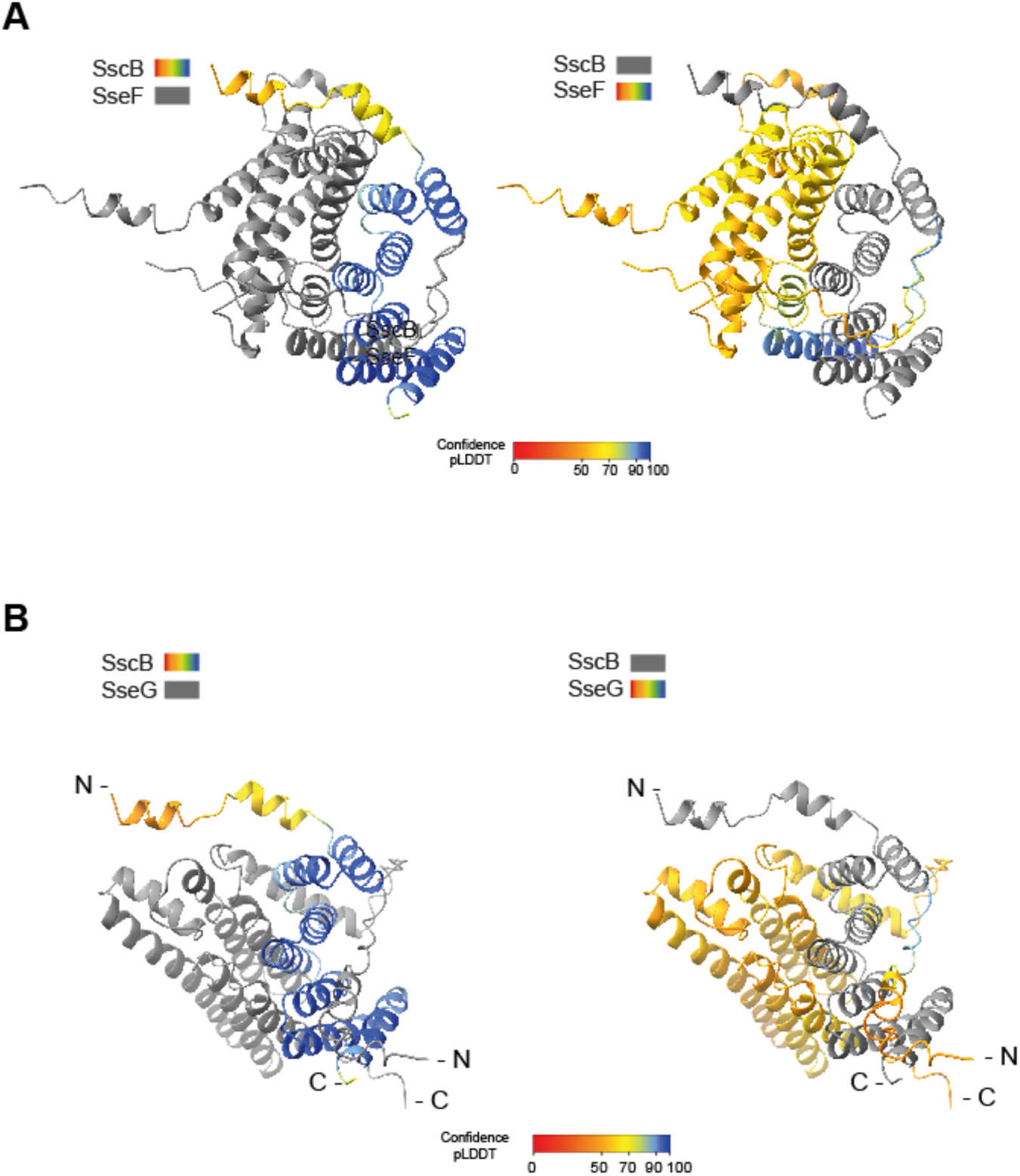
Alphafod2 multimer predictions of chaperone/effector complexes. **(A)** Predicted structure of SscB/SseF complex using Alphafold2 multimer. On the left, the coloured structure shows SscB in the complex. On the right, the coloured structure shows SseF in the complex. **(B)** Predicted structure of SscB/SseG complex using Alphafold2 multimer. On the left, the coloured structure shows SscB in the complex. On the right, the coloured structure shows SseG in the complex. The colour gradient indicates the confidence of the prediction (pLDDT).

**Figure S7:**
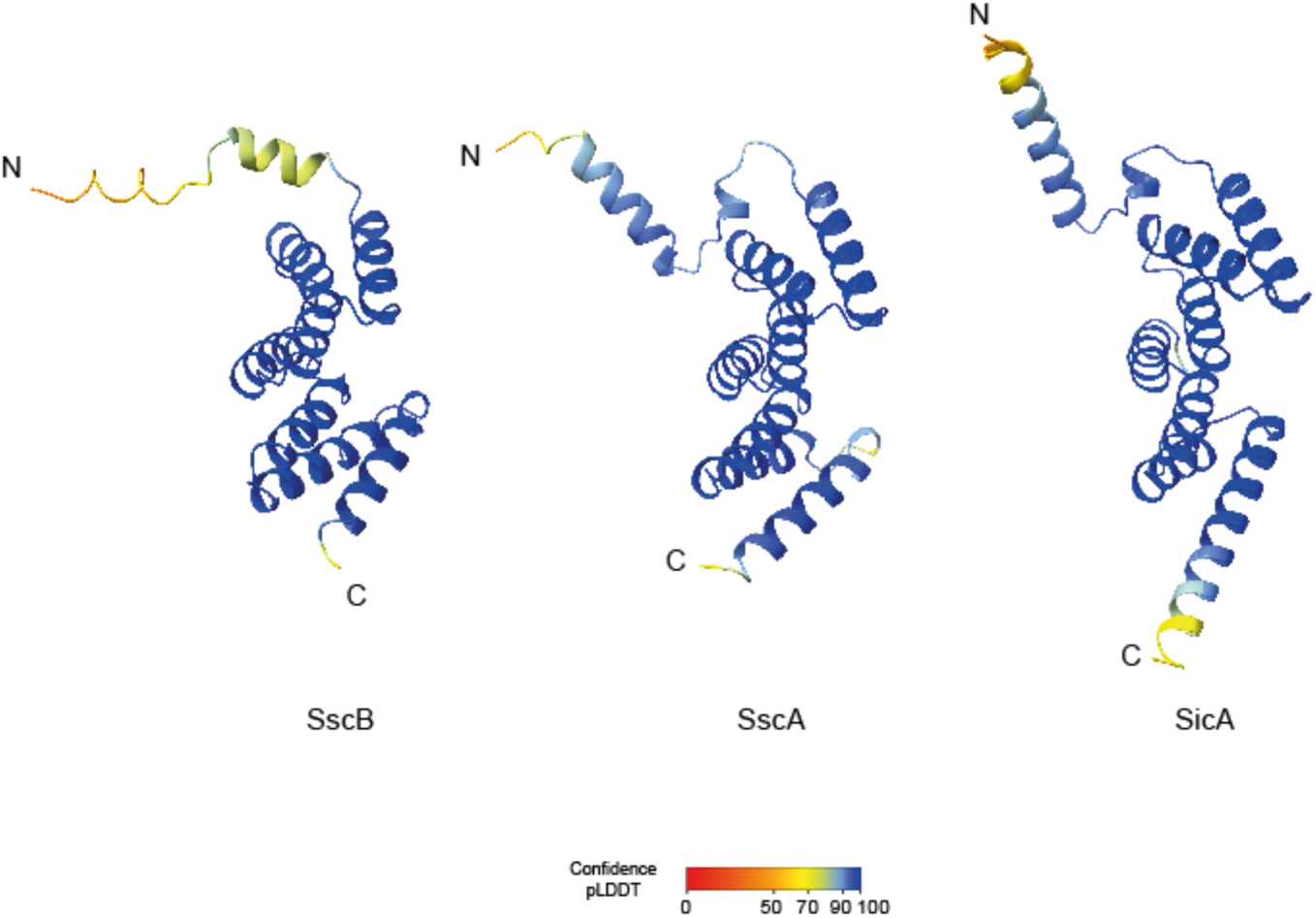
Alphafold2 predictions of T3S chaperone monomers. Predicted structure of monomeric SscB, SscA and SicA of *Salmonella* Typhimurium using Alphafold2. The colour gradient indicates the confidence of the prediction (pLDDT).

**Figure S8:**
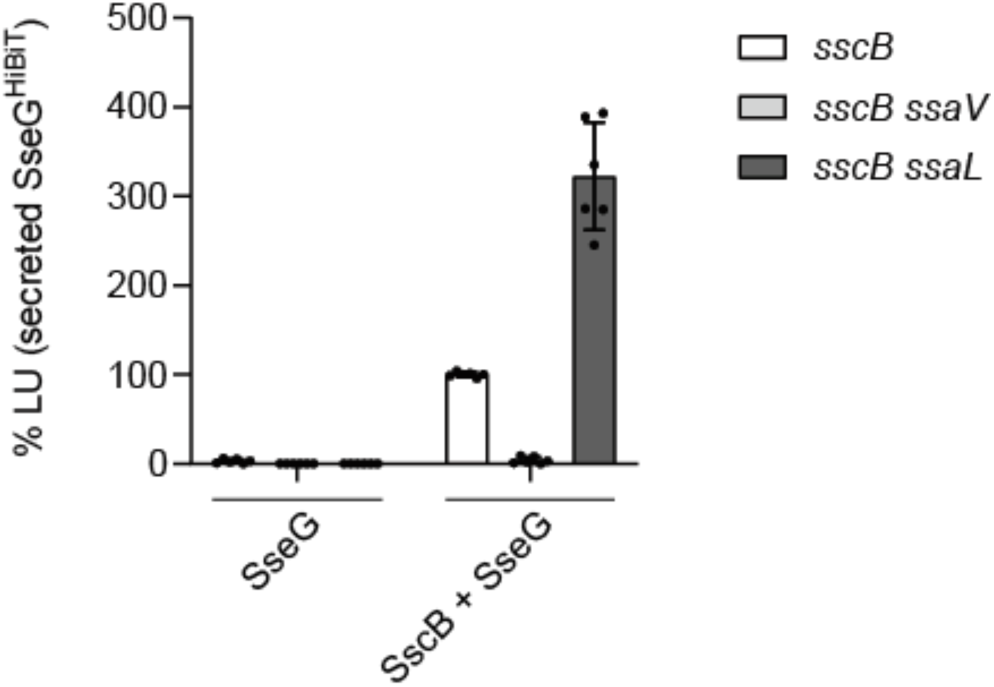
Secretion of SseG in different *Salmonella* mutant strains. T3S-proficient (*sscB*), T3S-deficient (*sscB ssaV*) and gatekeeper mutant (*sscB ssaL*) *Salmonella* strains, harbouring plasmids co-expressing SscB and SseG, under a rhamnose-inducible promoter, were grown under SPI-2-inducing and secreting conditions. Bacterial cultures were collected, and secretion was analysed using the split-NanoLuc T3S assay. The secretion of plasmid-derived SseG^HiBiT^ was analysed by luminometry. The luminescence signal was compared to the T3S-proficient strain (*sscB*) expressing SscB and SseG (SscB + SseG).

